# Learning to handle flowers increases pollen collection benefits for bees but does not affect pollination success for plants

**DOI:** 10.1101/2024.05.10.593657

**Authors:** Maggie M. Mayberry, Katherine C. Naumer, Annaliese N. Novinger, Dalton M. McCart, Rachel V. Wilkins, Haley Muse, Tia-Lynn Ashman, Avery L. Russell

## Abstract

Behavior frequently affects cooperation as well as conflict in plant-pollinator interactions. Pollinators such as bees often modify how they handle flowers with experience and such learning is generally assumed to increase collection of floral food rewards. The complexity of flower morphology also affects how quickly pollinators learn and recall how to access floral rewards from a given flower type. Because learning to handle a flower can increase extraction of food rewards (such as pollen) and often involves the pollinator altering how it interacts with the flower’s reproductive organs, pollinator learning could affect pollination success. Yet how pollinator cognition and flower morphology interact to affect pollination success is unknown. We therefore asked how learning and memory of flower handling by pollen-foraging generalist bumble bees (*Bombus impatiens*) varied among four morphologically distinct flower types *(Phacelia campanularia, Exacum affine*, *Solanum elaeagnifolium*, and *Erythranthe guttata*) and affected pollen collection and pollen deposition on these flowers. We found that bees learned and remembered how to handle some flower types more quickly than others. Learning typically involved development of motor routines unique to each flower type. While bees invariably learned to improve pollen collection, how quickly bees learned and remembered each flower type did not affect pollen collection. Surprisingly, pollen deposition on flowers was not affected by bee foraging experience. Thus, even though learning benefits the bee, it does not alter female (and potentially male) fitness benefits for the plant. We discuss potential reasons for these patterns and consequences for bee behavior and flower evolution.

## INTRODUCTION

Behavioral plasticity can substantially alter the degree to which interspecies interactions are mutually beneficial (Bronstein 2009; van der Kooi et al. 2021; Moran et al. 2022). Plant-pollinator mutualisms typically involve animals visiting flowers to collect food rewards (e.g., nectar and pollen), and in the process, picking up and transferring pollen to conspecific flowers. Because foraging can be dangerous and is expensive in terms of time and energy, pollinators are expected to optimize their foraging efficiency with experience (Lewis 1993; Laverty 1994; Chittka et al. 1999; Raine and Chittka 2008). For instance, over dozens or hundreds of flower visits, nectar foraging bees learn to make quicker visits and find nectar more quickly, which is predicted to increase nectar collection rate (Laverty 1994; Keasar et al. 1996; Evans et al. 2017; Barker et al. 2018). While learning is widely accepted to benefit the pollinator, learning could also affect benefits for the plant (Lewis 1993; Barker et al. 2018). Pollinator learning is traditionally thought to increase pollination success, for instance by increasing the likelihood that the pollinator stays true to one flower type (‘floral constancy’), thereby potentially reducing pollen loss and transfer to heterospecifics (Waser 1986; Chittka et al. 1999). Yet pollinator learning could also reduce pollination success. For example, experienced pollinators tend to make shorter visits that potentially result in fewer opportunities to pick up and transfer pollen (e.g., Laverty 1980; Gegear and Laverty 1995; Ramos et al. 2017). While learning is frequently hypothesized to change how the interests of plant and pollinator are aligned (e.g., Leonard and Papaj 2011; Evans et al. 2017; Barker et al. 2018; van der Kooi et al. 2021; Richman et al. 2021), evidence of how pollinator learning directly affects pollination success is extremely limited (see Internicola and Harder 2011; Vázquez and Barradas 2017; Ramos et al. 2017; Jones and Agrawal 2017).

Pollinator cognition is particularly well understood in the context of bee-flower interactions (Chittka and Thomson 2005; Giurfa 2007). While numerous studies over the past century have addressed learning and memory of generalist bees foraging for carbohydrate-rich nectar, bees must also collect protein-rich pollen to survive (Simpson and Neff 1981; Kevan and Baker 1983; Kitaoka and Nieh 2009; Nicolson and van Wyk 2011). Potentially hundreds of thousands of plant species offer pollen as a food reward to bees (Vogel 1978; Buchmann 1983; Russell et al. 2024), but we lack knowledge of whether pollen foragers, like nectar foragers, generally learn to optimize their flower handling and whether such learning results in increased pollen collection (see Raine and Chittka 2007; Russell et al. 2016). Nectar foragers often require dozens of floral visits to become proficient at handling flowers, particularly when flowers require complex motor routines (‘flower handling skills / techniques’) for bees to efficiently access concealed nectar (Laverty and Plowright 1988; Laverty 1994; Woodward and Laverty 1992; Chittka et al. 1999). Because flowers that offer pollen rewards also frequently possess diverse morphologies and often conceal their pollen (Russell et al. 2017; Table 1), it is reasonable to assume that pollen foragers should also have to learn to become proficient at handling flowers (Raine and Chittka 2007; Russell et al. 2016, 2017). Furthermore, while prior studies have focused on changes in time spent handling the flower, little quantitative work has examined how pollen foraging motor routines are modified with experience (see Laverty 1980; Chittka and Thomson 1997; Russell et al. 2016).

**Table 1:** Prominent features of the four kinds of pollen offering flowers.

| Flower | Pistil | Anther | Flower Morphology |
| --- | --- | --- | --- |
| *Phacelia campanularia* | Long, slender style with bilobed long and thin stigma | Five small anthers dehiscing completely with exposed pollen | Saucer shaped flower with anthers and style readily accessed |
| *Exacum affine* | Small stigma at the end of the single short style | Five short rigid anthers dehiscing by pores (pollen concealed) | Stellate flower with anthers and style readily accessed |
| *Solanum elaeagnifolium* | Small stigma at the end of a long style | Five long semi-rigid anthers dehiscing by pores (pollen concealed) | Stellate flower with anthers and style readily accessed |
| *Mimulus guttatus* | Long, slender style with bilobed sensitive stigma (closes with pollination) | Four very small anthers dehiscing completely with exposed pollen | Bell shaped flower with anthers hidden within corolla tube and style sometimes protruding |

Floral morphology often dictates how rewards are partitioned and dispensed and flower morphology frequently differs dramatically among plant species. As a result, efficient collection of rewards is hypothesized to require diverse flower handling techniques (Laverty and Plowright 1988; Lewis 1993; Gegear and Laverty 1995). For instance, foraging for nectar from a lupine flower (*Lupinus* spp.) requires the bee to force open the banner from the keel petals to extend its proboscis into the nectary (e.g., Westerkamp 1999), whereas bees must fully enter into the hood petals of wolfsbane flowers (*Aconitum* spp.) to locate the hidden nectaries (e.g., Laverty 1980; Laverty and Plowright 1988). Furthermore, the amount of learning required to become a proficient forager is hypothesized to depend on flower morphology, with relatively more learning required on flowers that require complex handling skills (Laverty 1980, 1994; Muth et al. 2015). Cognitive constraints associated with remembering complex flower handling techniques may result in this memory decaying more quickly than for simpler handling techniques (Chittka et al. 1999). Yet despite abundant evidence for how floral morphology affects learning and memory of nectar foraging bees, whether similar cognitive constraints are at play during pollen foraging have been barely explored (Raine and Chittka 2007; Russell et al. 2016).

How a flower’s morphology affects bee learning and memory may also affect benefits to both bee and plant (Vázquez and Barradas 2017; Ramos et al. 2017). For example, when less learning is required to become proficient at handling a given flower type, we expect bees to collect rewards sooner at their maximal rate. Likewise, relearning a handling technique should reduce pollen collection, but this reduction should be smaller when it is easier for the pollinator to remember how to handle the flower type. However, pollinator learning could be beneficial or costly for the plant (Table 2). For example, from the plant’s perspective learning might result in a better pollinator, if optimizing flower handling involves the bee contacting the plant reproductive organs more frequently or reliably. Yet learning could result in a worse pollinator from the plant’s perspective, if optimizing flower handling involves body positioning that circumvents contact with reproductive organs or making quicker visits that result in less pollen overall transferred among flowers. Furthermore, the more pollen moved into the pollen baskets (largely unavailable for pollination; Parker et al. 2015), the less pollen available for transfer among conspecific flowers (Castellanos et al. 2006; Hargreaves et al. 2009; Wilkins et al. 2022), which could further compound the negative effects of learning on pollination for the plant.

**Table 2:** Hypothesized benefits or costs of learning for bee and flower of measured attributes.

| Changes in | Potential consequences for bee | Potential consequences for plant |
| --- | --- | --- |
| **flower handling** |  |  |
| Reduced handling time with experience | Increases pollen collection rate as the bee becomes more efficient at handling the flower | Increases pollen deposition rate on flower stigmas as the bee visits flowers at a faster rate |
|  | Decreases pollen removal per flower as the bee optimizes time spent foraging on each flower | Decreases pollen deposition on flower stigmas if the bee spends less time contacting flower stigmas |
| Reduced latency to begin collecting pollen with experience | Increases pollen collection rate as the bee becomes quicker at locating the pollen reward | Increases pollen deposition on flower stigmas if the bee acquires pollen on its body earlier into flower visits. |
|  | N/A | Decreases pollen deposition on flower stigmas if the bee spends less time contacting stigmas due to finding the anthers earlier |
| Change in handling skills with experience | Increases pollen collection rate as the bee optimizes its flower handling skills | Increases pollen acquisition and transfer if the flower anthers and stigma are contacted more effectively |
|  | Decreases pollen removal per flower as the bee optimizes how much effort to expand on each flower | Decreases pollen deposition on flower stigmas if the bee becomes more effective at collecting pollen overall |

In this laboratory study, we assessed how diverse floral morphologies influenced learning and memory of flower handling by pollen foraging bumble bees (*Bombus impatiens*), and how acquisition of these skills influenced pollen collection and pollen transfer among flowers. We used fresh flowers of four nectarless species (*Phacelia campanularia* (Boraginaceae), *Exacum affine* (Gentianaceae), *Solanum elaeagnifolium* (Solanaceae), and *Erythranthe guttata* (Phrymaceae)) that offer only pollen as a reward, but which differ substantially in their morphology and in how pollen is dispensed (Figure 1; Table 1). We hypothesized that 1) flower types differ in handling difficulty and in how readily handling skills are learnt and remembered, and that 2) these differences affect how much pollen collection increases with foraging experience. To be precise, we expected that differences in pollen collection between flower-naïve and experienced bees would be greatest when learning was more difficult. We also hypothesized that 3) differences in learning and memory of flower handling would affect pollination success from the plant’s perspective. Specifically, we expected that pollen transfer would decline as bees gained experience and decline faster when learning was easier.

**Figure 1.**
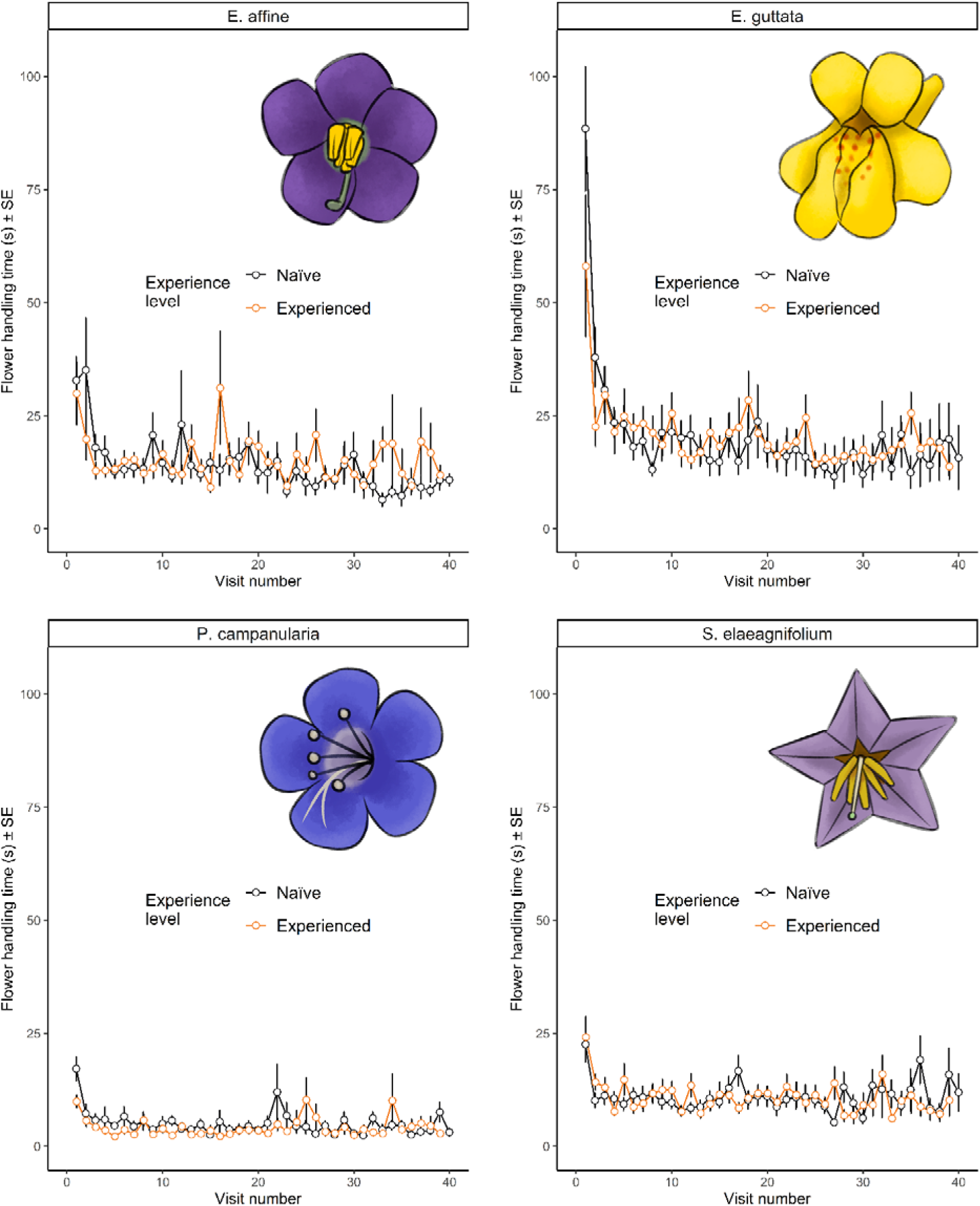
Handling times shown for the first 40 (out of up to 70) flower visits made by initially naïve and experienced bees on each plant species. *N* = 21, 20, 17, 20 initially naïve bees and *N* = 21, 20, 19, 20 experienced bees foraging on *E. affine*, *E. guttata, P. campanularia*, and *S. elaeagnifolium*, respectively.

## METHODS

### Experimental subjects

To study how pollen foraging experience on different flowers affected flower handling skills, pollen collection, and pollen transfer, we used 81 initially flower-naïve workers from seven captive commercially obtained colonies (Koppert Biological Systems, Howell, MI, USA) of the common eastern bumble bee, *Bombus impatiens*. Briefly, following Russell et al. 2017, each colony was maintained on 2 molar solution of sucrose and pulverized honeybee-collected pollen (Koppert Biological Systems) from artificial feeders within enclosed foraging arenas (LWH: 82 × 60 × 60 cm) set to a 14 h:10 h light : dark cycle.

We used fresh flowers from four plant species differing substantially in floral morphology (Table 1; Figure 1) as donors and recipients of conspecific pollen. Following Laverty (1994) and Laverty and Plowright (1988), we reasoned that differences in floral morphology would correspond to differences in learning and memory required to improve extraction of resources. Flowers used in trials were freshly cut from 10 *Phacelia campanularia* (Boraginaceae), 36 *Exacum affine* (Gentianaceae), 8 *Solanum elaeagnifolium* (Solanaceae), and 10 *Erythranthe guttata* (Phrymaceae) plants, which were grown in a university greenhouse with supplemental halogen lights to extend day length to a 14:10 h light : dark cycle, and were fertilized every other week (PlantTone, NPK 5:3:3, Espoma, Millville, New Jersey, USA). To prevent desiccation, all freshly cut live flowers were placed into custom water tubes (Russell et al. 2017). All flowers used offer only pollen as a reward; they do not produce nectar.

### Experimental protocol

To identify appropriate test subjects, we captured bees observed foraging on the artificial nectar feeders, painted them with unique color combinations using non-toxic oil markers (Sharpie, CA), and returned them to their colonies. We divided painted flower-naïve bees into four treatments, with two sub-treatments each. A minimum of three colonies were represented per treatment. Treatments differed in terms of the plant species that was used: 20 cut flowers of a given plant species were spaced 7 cm apart in a 5 x 4 Cartesian grid design on the arena wall. We systematically alternated assignment of bees to each treatment to control for effects of time and day on behavior. Sub-treatments differed in terms of whether a bee was initially flower-naïve (‘naïve’) or had previously been tested (‘experienced’): each bee therefore received two consecutive trials with the same plant species, with the naïve and experienced trial separated by approximately 24 hours. After its naïve trial, the bee was returned to its colony, but after its experienced trial, the bee was euthanized by freezing in a -20 °C freezer, to facilitate other workers to forage.

To initiate a behavioral trial, flowers were set up and a single painted flower-naïve worker bee was gently captured from the foraging arena using a 40 dram vial (Bioquip) and immediately released in the center of the test arena following Russell et al. (2017). Before releasing a bee, we visually confirmed the absence of pollen on her body. To estimate how much experience was required to reach an asymptote in flower handling skill, we ran several long practice trials (120 visits) and estimated that an asymptote was reached less than 50% through each practice trial; data from these practice trials was not used in any subsequent analyses. We therefore terminated each naïve behavioral trial after 70 visits (or earlier if the bee stopped visiting flowers for 5 minutes) to ensure that bees had learned and to avoid bees depleting flowers of pollen rewards or filling their pollen baskets completely. For a subset of flowers and trials we extracted pollen from the anthers to confirm that pollen had not been depleted after a trial. We terminated each experienced behavioral trial immediately once the bee had made the same number of visits as in its naïve trial. To terminate a trial, we turned off the overhead arena lights and captured the bee in a vial. After terminating a trial, the complete pollen load from one pollen basket (i.e., corbicular pollen load) was carefully removed from the bee, and the anthers and styles of all visited flowers were removed. The pollen load, anthers, and styles were stored separately in 70% ethanol for pollen counting (styles and anthers for each trial were separately pooled into microcentrifuge tubes). The arena was cleaned thoroughly between trials.

Trials were video recorded to enable using an event logging program (BORIS v.8.19.3) to measure the handling time per flower visit (from three legs on the flower until physical contact with the flower ended), the latency to begin collecting pollen after landing on the flower (from three legs on the flower until the bee began scrabbling or buzzing on the anthers), and the duration of each flower visit during which bees used easily distinguishable categories of pollen collecting handling routines (Figure 5; see Results for a description). We also recorded whether the bee had collected pollen on a given flower visit. The naïve trial for three bees failed to be video recorded and this data was therefore unavailable for behavioral analyses.

### Assessment of pollen collection and transfer

To examine whether bee foraging experience affected pollen collection, the corbicular pollen load from each trial was stored in separate microcentrifuge tubes with 1000 μL of 70% ethanol. We counted pollen in two or three 10 μL aliquots using a haemocytometer (Hausser Scientific, Horsham, PA) at 400× or 100× (Leica DM 500) to arrive at an estimate for the total volume. Estimated pollen counts were rounded to the nearest whole number. To evaluate pollen deposition, the pollen grains on flower stigmas was enumerated. We acetolyzed styles from flowers (following Dafni 1992) pooled by trial, condensed samples by centrifugation to 40 μL, and counted resuspended pollen in two 10 μL aliquots; if we counted zero grains, we counted conspecific grains in all four aliquots.

### Data Analyses

All data were analyzed using R v.4.2.2 (R Development Core Team 2023).

#### Does foraging experience increase efficiency to handle flowers and begin collecting pollen and do patterns depend on plant species?

We analyzed how foraging experience affected the relative efficiency of naïve bees in handling flowers and in their latency to begin collecting pollen after landing on a flower. Following Laverty (1994) and Laverty and Plowright (1988), we assumed that experienced bees had achieved maximal efficiency and that naïve bees increased their efficiency with foraging experience. We therefore calculated the relative efficiency of a given naïve bee as a percentage of its maximum efficiency (the mean handling time or latency during its experienced trial). For all naïve bees we thus calculated initial foraging efficiency for the first flower visit, as well as the rate of improvement in efficiency (slope from linear regressions) until reaching an asymptote in performance (three consecutive visits with ≥ 90% efficiency; following Laverty 1994). For bees that did not reach an asymptote in efficiency for flower handling (3 of 78 bees) or latency to begin collecting pollen (10 of 78 bees), all visits were used in regressions following Laverty (1994). We used Kruskal-Wallis Tests (KWT) to assess whether the initial efficiency, slope, or number of visits until reaching the asymptote differed among plant species, and in cases of significance we used a Wilcoxon-Signed Rank Test (WSRT) with corrections for multiple testing to determine pairwise differences. The fit of the regressions was good overall (coefficient of determination mean ± SE: handling time: *P. campanularia*: 0.61 ± 0.09; *E. affine*: 0.51 ± 0.09; *S. elaeagnifolium*: naïve: 0.51 ± 0.09; *E. guttata*: 0.57 ± 0.07; latency to begin collecting pollen: *P. campanularia*: 0.42 ± 0.10; *E. affine*: 0.38 ± 0.07; *S. elaeagnifolium*: 0.32 ± 0.7; *E. guttata*: 0.59 ± 0.07).

#### Does memory of improvements in flower handling and latency to collect pollen differ among plant species treatments?

To analyze how memory (e.g. overnight retention) of improvements in handling time or latency to begin collecting pollen after landing differed among bees foraging on different plant species, we applied a learning curve to the first 20 floral visits of each naïve and experienced bee and analyzed the estimated parameters. We chose this standardized cutoff because most naïve bees reached an asymptote in efficiency (see above) within 20 visits (96% and 85%, for handling time and latency to collect pollen, respectively) and because rates of improvement in efficiency did not differ significantly among plant species (see Results). We used a Wright’s cumulative average model (Martin, n.d.). The model takes the form Y = *a*X*^b^*, where *Y* is the cumulative handling time or latency to collect pollen (measured in seconds) per floral visit, *X* is the cumulative number of floral visits, *a* is the estimated handling time or latency to collect pollen for the first floral visit (intercept), and *b* is the slope of the function in log-log space.

The fit of the model was good overall: the mean coefficient of determination was high (mean ± SE: handling time: *P. campanularia*: naïve: 0.74 ± 0.08; experienced: 0.77 ± 0.06; *E. affine*: naïve: 0.74 ± 0.06; experienced: 0.63 ± 0.05; *S. elaeagnifolium*: naïve: 0.68 ± 0.07; experienced: 0.67 ± 0.07; *E. guttata*: naïve: 0.84 ± 0.05; experienced: 0.74 ± 0.07; latency to begin collecting pollen: *P. campanularia*: naïve: 0.56 ± 0.06; experienced: 0.58 ± 0.06; *E. affine*: naïve: 0.74 ± 0.06; experienced: 0.65 ± 0.06; *S. elaeagnifolium*: naïve: 0.59 ± 0.06; experienced: 0.76 ± 0.06; *E. guttata*: naïve: 0.81 ± 0.05; experienced: 0.70 ± 0.08).

We then analyzed these parameters using generalized linear mixed effects models (GLMM) with a gaussian distribution using the glmer() function in the lme4 package (Bates et al. 2015), specifying type II Wald Chi-square (χ2) tests via the Anova() function in the car package (Fox 2015). The response variable was the slope of the learning curve (parameter *b*) or the intercept (parameter *a*). We deleted three measurements identified as outliers by the plot_model() function (sjPlot package; Lüdecke et al. 2021). We specified explanatory variables for all GLMMs used in this study as ‘bee experience level’ (naïve or experienced) and ‘plant species’ (*P. campanularia*, *E. affine*, *S. elaeagnifolium*, or *E. guttata*) and nested random effects for all GLMMs as ‘bee ID’ within ‘colony ID’. We checked for overdispersion, zero inflation, and uniformity for all GLMMs using the DHARMa package (Hartig 2018) and log transformed pollen counts and total handling time to meet model assumptions.

#### Do flower handling skills change with experience and depend on plant species?

To analyze how flower handling skills changed with bee foraging experience, we used GLMMs to assess whether bees were more likely to access pollen with experience and whether handling routines used to collect pollen changed with experience, with explanatory variables as above. For the first GLMM, we specified the response variable as ‘visits to 80% learning criterion’ (number of visits required to successfully collect pollen on 8 of the last 10 flower visits), log transformed to meet model assumptions. For the second GLMM, we specified the response variable as ‘mean percent modified handling routine’ (percent of total handling time for a flower that the bee used the given modified handling routine, averaged across all flower visits for the given trial). To meet model assumptions, we added 1% to this response variable and square root transformed it.

#### Does pollen collection increase with experience and do patterns depend on plant species?

To analyze how the quantity and rate of pollen grains collected into bee pollen baskets was influenced by foraging experience, we first normalized pollen collection by the number of successful pollen collecting flower visits (i.e., for each trial: total pollen collected divided by total pollen collecting visits made). We then ran GLMMs with ‘mean pollen collected or ‘mean pollen collection rate’ (mean pollen collected divided by total time spent handling flowers) as the response variable, with explanatory variables as above.

#### Does foraging experience affect pollen receipt by stigmas and do patterns depend on plant species?

To analyze how the quantity of pollen grains bees deposited on flower stigmas was influenced by bee foraging experience, we first normalized pollen deposition by the number of flower visits made (i.e., for each trial: total pollen deposition divided by total visits made, regardless of successful pollen collection). We then ran GLMMs with ‘mean pollen deposited’ or ‘mean pollen deposition rate’ (mean pollen deposited divided by total time spent handling flowers) as the response variable, with explanatory variables as above.

## RESULTS

### Efficiency to handle flowers and begin collecting pollen increased with experience, but differed little among plant species

Naïve bees differed significantly in their initial flower handling efficiency among the four different plant species (Figure 1a; KWT: χ^2^_3_ = 14.54, *P* < 0.003). Specifically, bees visiting *E. guttata* and *P. campanularia* had a significantly lower initial handling efficiency than bees visiting either *E. affine* or *S. elaeagnifolium* (WSRT: *P* < 0.02; mean % ± SE efficiency relative to experienced state: for *P. campanularia*: 40 ± 10; for *E. guttata*: 41 ± 9; for *E. affine*: 74 ± 12; for *S. elaeagnifolium*: 101 ± 22). However, this did not translate to a significant difference among plant species in the number of flower visits required to reach an asymptote in handling efficiency (KWT: χ^2^_3_ = 3.99, *P* = 0.263). Across all treatments, naïve bees rapidly learned within 6 to 9 flower visits on average (mean visits ± SE: for *E. affine*: 5.90 ± 0.96; for *S. elaeagnifolium*: 5.95 ± 1.44; for *P. campanularia*: 7.41 ± 1.6; for *E. guttata*: 8.45 ± 1.42). Accordingly, naïve bees did not differ significantly in their rate of improvement in flower handling efficiency (KWT: χ^2^_3_ = 0.34, *P* = 0.953). On average, efficiency per visit increased quickly, by 30%, 32%, 43%, and 70% on *E. guttata*, *E. affine*, *S. elaeagnifolium*, and *P. campanularia*, respectively.

In addition, neither the initial efficiency in latency to extract pollen after landing (mean % ± SE efficiency relative to experienced state: for *E. guttata*: 54.9 ± 12.9; for *S. elaeagnifolium*: 74 ± 13; for *E. affine*: 98 ± 23; for *P. campanularia*: 99 ± 23), nor the rate of improvement (efficiency per visit increased by 11%, 19%, 20%, and 42% on *S. elaeagnifolium*, *E. guttata*, *E. affine*, and *P. campanularia*, respectively) differed among naïve bees visiting different plant species (Figure 1b; KWT: initial efficiency: χ^2^_3_ = 4.07, *P* = 0.254; rate of improvement: χ^2^_3_ = 5.708, *P* = 0.127). There was also no significant difference in the number of flower visits required to reach an asymptote in performance among plant species (KWT: χ^2^_3_ = 0.027, *P* = 0.999). On average, naïve bees quickly learned within 9 to 15 flower visits (mean visits ± SE: for *E. guttata*: 9.05 ± 1.64; for *E. affine*: 9.5 ± 2.02; for *S. elaeagnifolium*: 11.20 ± 2.70; for *P. campanularia*: 14.71 ± 4.80).

### How well bees remembered improvements in flower handling and latency to collect pollen depended on plant species

While bees rapidly learned to handle flowers (Figure 1) and begin collecting pollen (data not shown), memory was imperfect (Figure 1, 2a, 3a). The degree to which memory decayed overnight depended significantly on plant species, with the most relearning required when foraging on *S. elaeagnifolium* and the least on *E. guttata* (Figure 1, 2a,b; learning rate GLMMs: handling time: effect of experience: χ^2^_1_ = 15.17, *P* < 0.0001; effect of plant species: χ^2^_3_ = 11.71, *P* < 0.009; effect of plant species X experience: χ^2^_3_ = 14.50, *P* < 0.003; Figure 1, 3a,b; latency to collect pollen: effect of experience: χ^2^_1_ = 1.20, *P* = 0.273; effect of plant species: χ^2^_3_ = 10.75, *P* < 0.014; effect of plant species X experience: χ^2^_3_ = 12.85, *P* < 0.005). Differences between naïve and experienced bees in handling time for the first flower (37%, 32%, and 22% mean decrease for *P. campanularia*, *E. guttata*, and *E. affine* flower, respectively, and a 12% mean increase for *S. elaeagnifolium*) revealed much the same pattern, with *P. campanularia* and *E. guttata* requiring less relearning (Figure 1, 2b; GLMM: effect of experience: χ^2^_1_ = 28.18, *P* < 0.0001; effect of plant species: χ^2^_3_ = 80.02, *P* < 0.0001; effect of plant species X experience: χ^2^_3_ = 3.66, *P* < 0.017). Differences between naïve and experienced bees in the latency to extract pollen from the anthers of the first flower (0.7%, 20%, and 46% mean decrease for *P. campanularia*, *E. affine*, and *E. guttata*, respectively, and a 4% mean increase for *S. elaeagnifolium*) also suggest that *E. guttata* required the least relearning (Figure 1, 3b; GLMM: effect of experience: χ^2^_1_ = 10.81, *P* < 0.002; effect of plant species: χ^2^_3_ = 140.70, *P* < 0.0001; effect of plant species X experience: χ^2^_3_ = 21.05, *P* < 0.0002).

**Figure 2.**
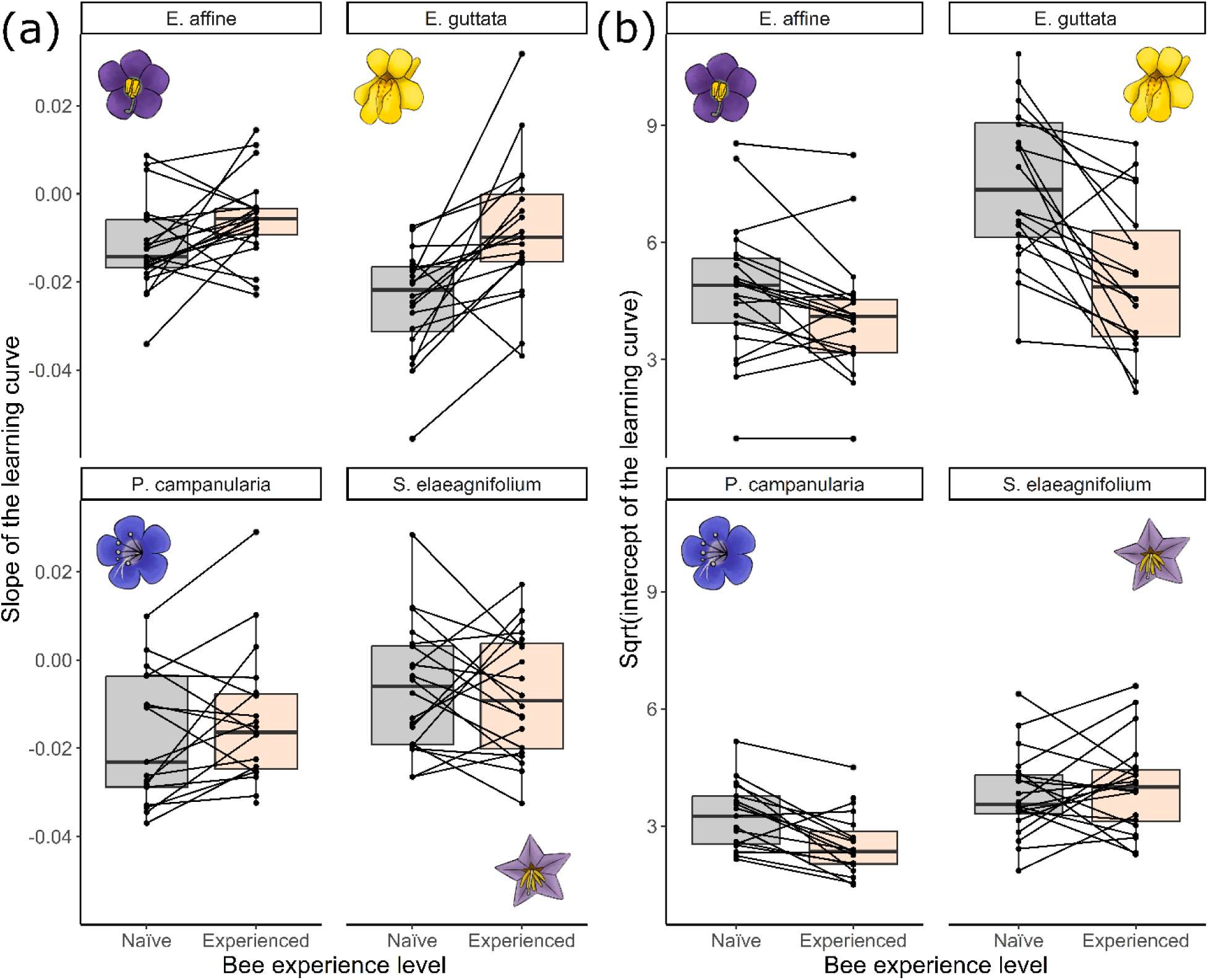
The predicted (a) rate of learning to handle flowers (slope of the learning curve) and (b) handling time of the first flower visit (intercept of the learning curve), as influenced by bee experience and plant species (same dataset as Figure 1). Plotted as boxplots with raw data indicated via dots joined by lines showing how individual bees altered their flower handling time with experience. *N* = 21, 20, 17, 20 initially naïve bees and *N* = 21, 20, 19, 20 experienced bees foraging on *E. affine*, *E. guttata, P. campanularia*, and *S. elaeagnifolium*, respectively.

**Figure 3.**
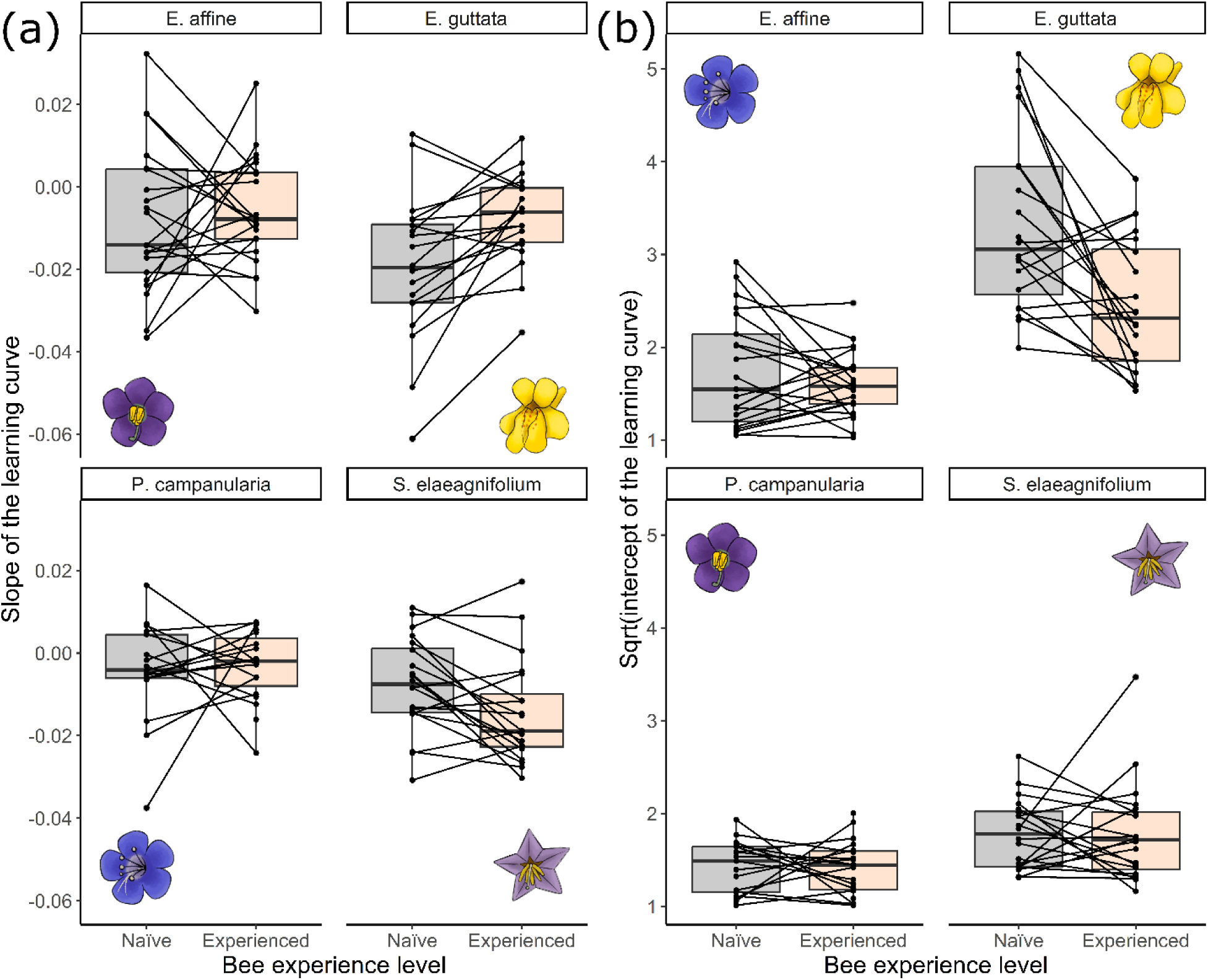
The predicted (a) rate of change in the latency to begin collecting pollen after landing (slope of the learning curve) and (b) latency to begin collecting pollen after landing on the first flower visit (intercept of the learning curve), as influenced by bee experience and plant species. Plotted as boxplots with raw data indicated via dots joined by lines showing how individual bees altered their latency to collect pollen with experience. *N* = 21, 20, 17, 20 initially naïve bees and *N* = 21, 20, 19, 20 experienced bees foraging on *E. affine*, *E. guttata, P. campanularia*, and *S. elaeagnifolium*, respectively.

### Flower handling skills changed with experience and depended on plant species

While all bees collected pollen from flowers during trials, bees did not always successfully collect pollen on a given floral visit. Across all bees, meeting the 80% learning criterion to successfully extract pollen (8 of the last 10 flower visits) took between 10 and 67 visits (Figure 4). Overall, naïve bees took more visits to meet the learning criterion than experienced bees, though this effect depended strongly on plant species (Figure 4; GLMM: effect of experience: χ^2^_1_ = 3.88, *P* < 0.049; effect of plant species: χ^2^_3_ = 58.9, *P* < 0.0001; effect of plant species × experience: χ^2^_3_ = 25.87, *P* < 0.0001). This effect was largely driven by bees foraging on *E. guttata*, which, after experience, took on average 73% fewer visits to meet the learning criterion (a further 4 of 20 naïve bees on *E. guttata* never met the criterion). Additionally, the proportion of naïve bees that failed to collect pollen on their very first visit differed significantly among plant species (on *S. elaeagnifolium*: 0%; on *P. campanularia*: 24%; on *E. affine*: 33%; on *E. guttata*: 55%; TEP: χ^2^_3_ = 15.404, *P* < 0.002).

**Figure 4.**
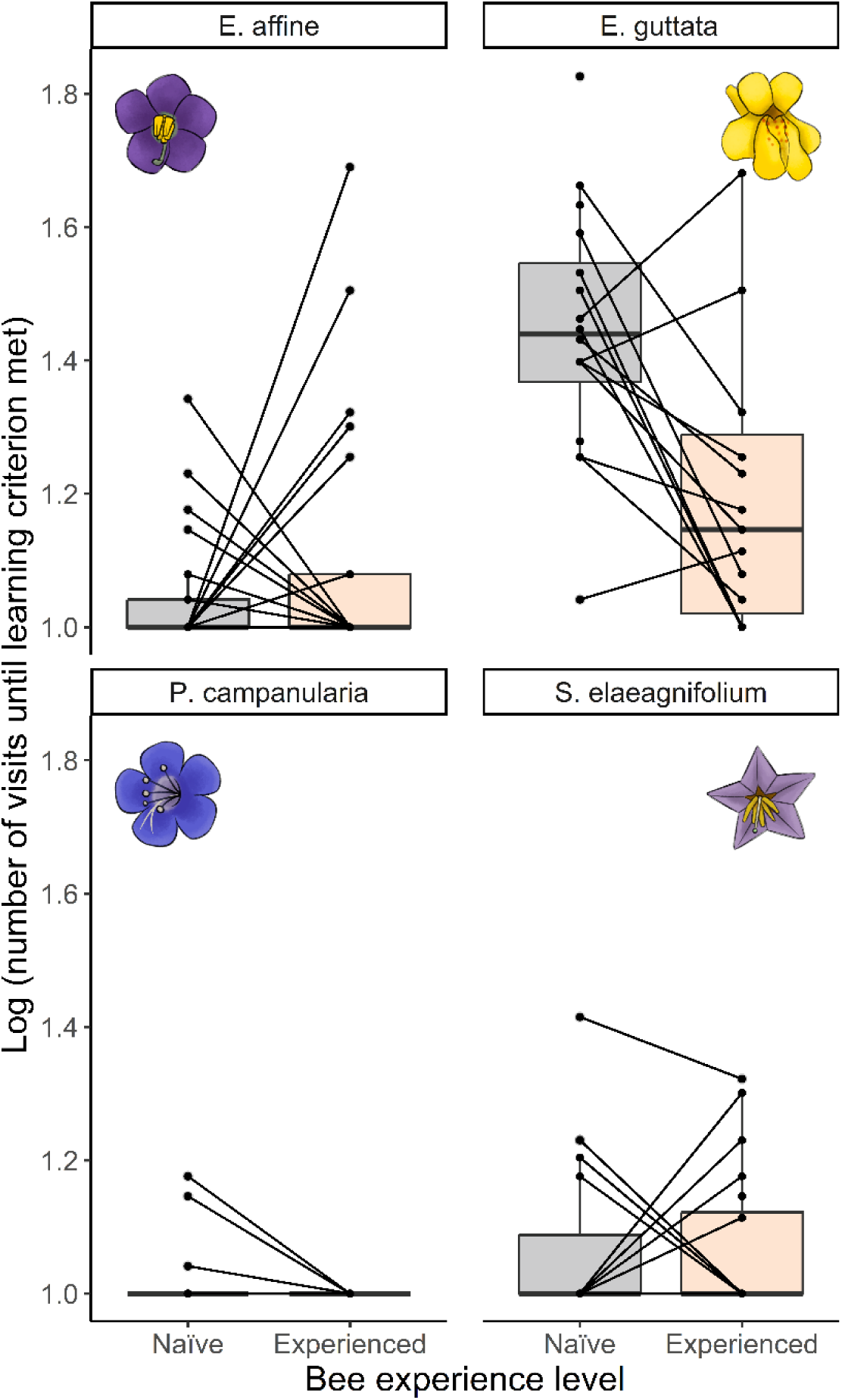
Number of flower visits required to reach the 80% learning criterion (successfully collecting pollen on 8 of the last 10 visits). Plotted as boxplots with raw data indicated via dots joined by lines showing how pollen collection success by individual bees changed with experience. *N* = 21, 20, 17, 20 initially naïve bees and *N* = 21, 20, 19, 20 experienced bees foraging on *E. affine*, *E. guttata, S. elaeagnifolium*, and *P. campanularia* respectively.

With experience, bees also significantly altered handling routines used to collect pollen and the magnitude of this change differed significantly among plant species (Figure 5a; GLMM: effect of experience: χ^2^_1_ = 24.82, *P* < 0.0001; effect of plant species: χ^2^_2_ = 99.64, *P* < 0.0001; effect of plant species × experience: χ^2^_2_ = 2.22, *P* = 0.330). Naïve bees on *E. affine* primarily bit down and buzzed the dorsal surface of the anthers to extract pollen on each visit (per visit, mean % ± SE: 79 ± 2.4), but with experience, bees often inverted themselves on the downward curving anthers, biting them and buzzing while anther pores were appressed to the ventral thorax (Figure 5b; 32 ± 2.9 % of the time). Buzzes, which indicated an attempt at extracting pollen, were identified by their distinctive sound and occurred only after a bee had landed (see Russell et al. 2017). On *S. elaeagnifolium*, naïve bees primarily bit and buzzed the individual spread apart anthers (53 ± 3.2 % of the time), while with experience, bees more commonly (53 ± 2.7 %) clasped three or more anthers together while biting and buzzing (Figure 5b). Naïve bees on *E. guttata* almost entirely scrabbled for pollen (using their legs and mandibles to knock pollen from the anthers; 95 ± 2.0 % of the time; Russell and Papaj 2016) while upright within the corolla tube. In contrast, as bees on *E. guttata* gained experience, they began inverting themselves within the corolla tube and buzzed pollen directly from the anthers (Figure 5b; 12 ± 3.5 % of the time). We were unable to similarly objectively characterize changes in the pollen foraging handling routines of bees on *P. campanularia*, but even on their very short visits to this flower, bees may have altered how they handled the anthers with experience. Altogether, after 24 hours, experienced bees had increased their use of the modified handling techniques on *S. elaeagnifolium*, *E. affine*, and *E. guttata* on average by 14%, 54%, and 127%, respectively.

**Figure 5.**
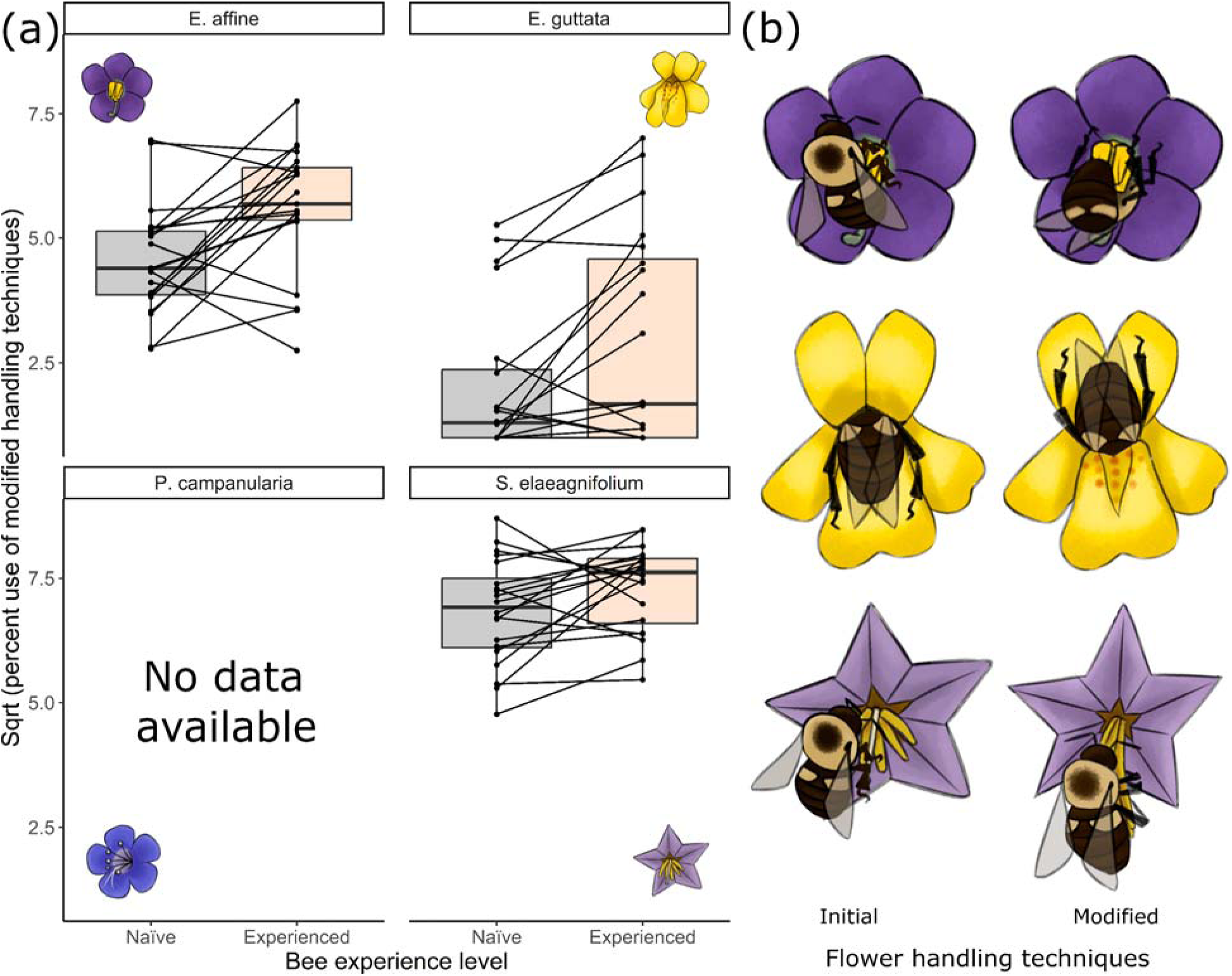
(a) Illustrations of the initial and modified handling techniques (‘motor routines’) used to extract pollen from flowers and (b) mean percent of a flower visit during which the modified handling routines were used to extract pollen. Plotted as boxplots with raw data indicated via dots joined by lines showing how handling routines used by individual bees changed with experience. No data are available for *P. campanularia* as we were unable to objectively characterize changes in the pollen foraging motor routine on this flower type. *N* = 21, 20, 20 initially naïve bees and *N* = 21, 20, 20 experienced bees foraging on *E. affine*, *E. guttata,* and *S. elaeagnifolium*, respectively.

### Bees collected more pollen more quickly into their pollen baskets with experience, but this pattern did not differ among plant species

The quantity of pollen grains bees collected into their pollen baskets and rate of collection increased significantly with experience and did not depend on plant species (Figure 6a; Table 3; GLMM for quantity: effect of experience: χ^2^_1_ = 13.26, *P* < 0.0003; effect of experience X plant species: χ^2^_3_ = 0.39, *P* = 0.943; Figure S1a, GLMM for rate: effect of experience: χ^2^_1_ = 9.88, *P* < 0.002; effect of experience × plant species: χ^2^_3_ = 0.92, *P* = 0.820). With experience, bees foraging on *P. campanularia*, *E. affine*, *S. elaeagnifolium*, and *E. guttata* collected on average 17%, 19%, 26%, and 27% more pollen, respectively. Predictably, given the large differences in the amount of pollen offered by the flowers of each plant species (following Table 3), the quantity and rate of pollen collection by bees differed significantly across plant species (Figure 6a; Table 3; GLMM for quantity: effect of plant species: χ^2^_3_ = 375.68, *P* < 0.0001; Figure S1a, GLMM for rate: effect of plant species: χ^2^_3_ = 181.06, *P* < 0.0001).

**Figure 6.**
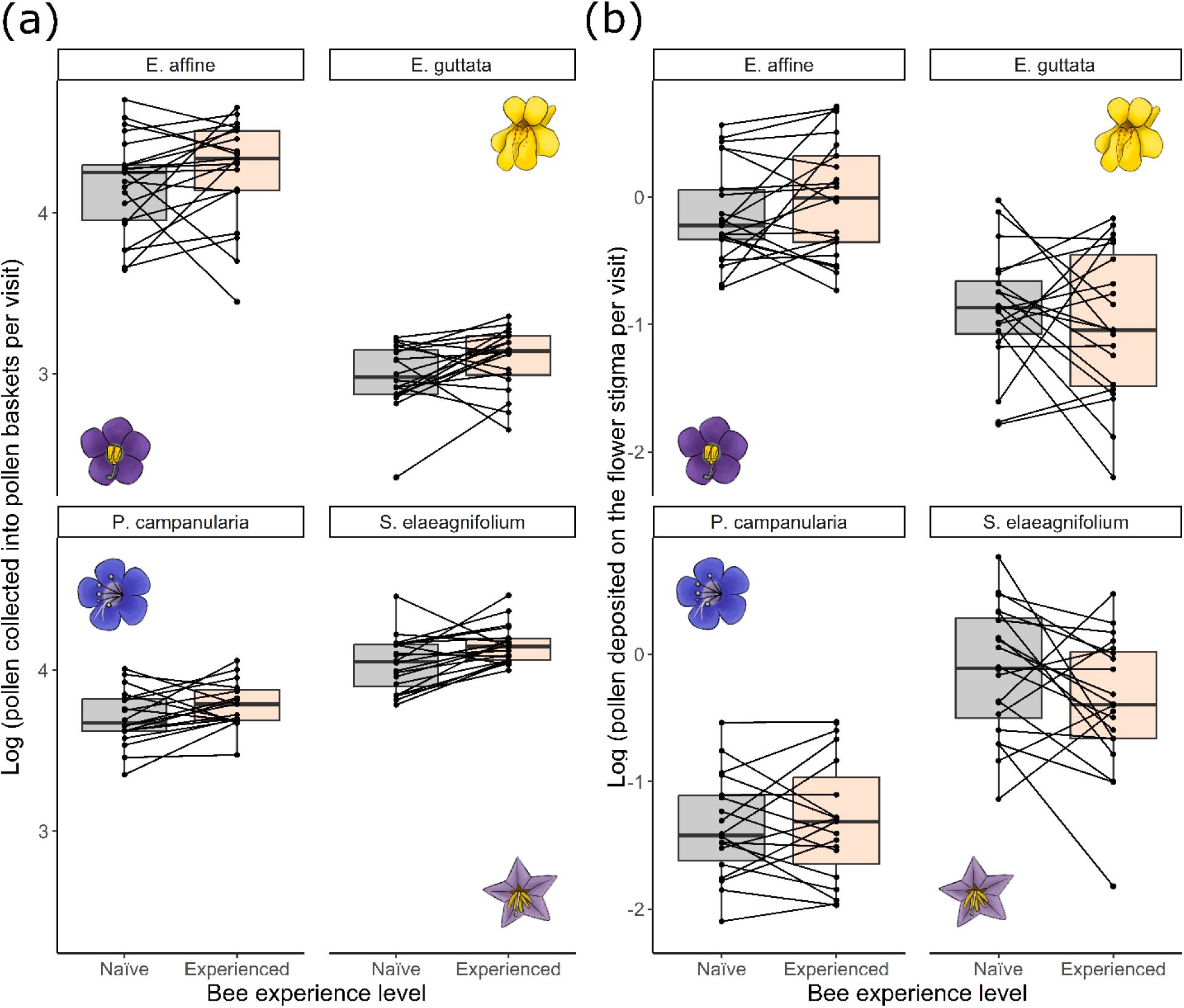
The quantity of pollen grains (a) in the pollen baskets (corbiculae) or (b) transferred to flower stigmas, as influenced by bee experience and plant species. Plotted as boxplots with raw data indicated via dots joined by lines showing how the amount of pollen collected or transferred by individual bees changed with experience. *N* = 21, 20, 17, 20 initially naïve bees and *N* = 21, 20, 19, 20 experienced bees foraging on *E. affine*, *E. guttata, P. campanularia*, and *S. elaeagnifolium*, respectively.

**Table 3:**
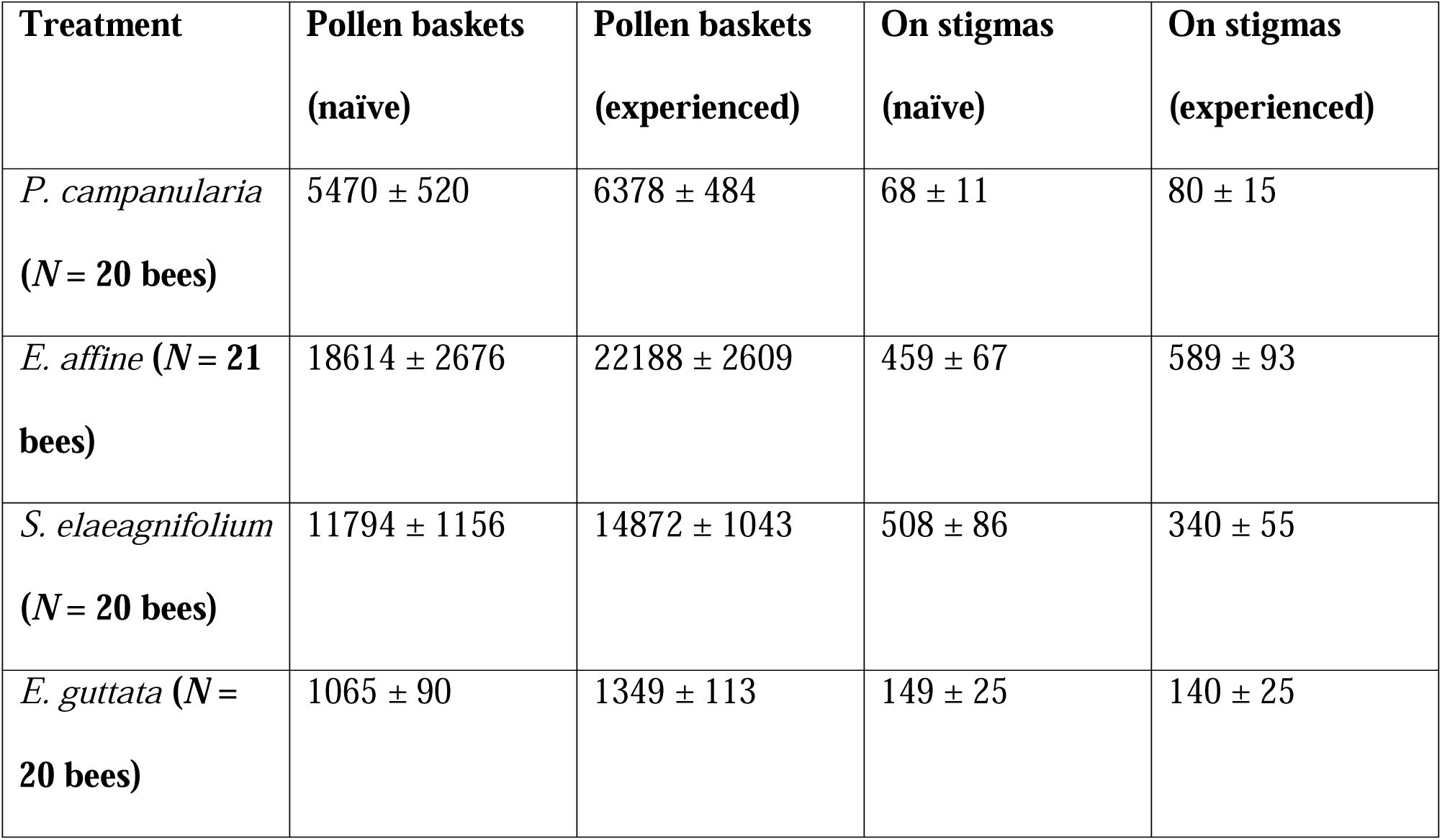
Quantity of pollen grains per flower visit (mean ± SE) collected by bees into the pollen baskets and transferred to stigmas among the flower treatments.

| **Treatment** | **Pollen baskets**<br>**(naïve)** | **Pollen baskets**<br>**(experienced)** | **On stigmas**<br>**(naïve)** | **On stigmas**<br>**(experienced)** |
| --- | --- | --- | --- | --- |
| *P. campanularia*<br>(*N* = 20 bees) | 5470 ± 520 | 6378 ± 484 | 68 ± 11 | 80 ± 15 |
| *E. affine* (*N* = 21<br>bees) | 18614 ± 2676 | 22188 ± 2609 | 459 ± 67 | 589 ± 93 |
| *S. elaeagnifolium*<br>(*N* = 20 bees) | 11794 ± 1156 | 14872 ± 1043 | 508 ± 86 | 340 ± 55 |
| *E. guttata* (*N* =<br>20 bees) | 1065 ± 90 | 1349 ± 113 | 149 ± 25 | 140 ± 25 |

### Bee foraging experience did not affect pollen receipt by stigmas for any plant species

Foraging experience did not affect the quantity of pollen grains deposited on flower stigmas by bees or the rate of pollen deposition, regardless of plant species (Figure 6b, Table 3; GLMM for quantity: effect of experience: χ^2^_1_ = 0.76, *P* = 0.384; effect of experience × plant species: χ^2^_3_ = 6.07, *P* = 0.108; Figure S1b, GLMM for rate: effect of experience: χ^2^_1_ = 0.52, *P* = 0.469; effect of experience × plant species: χ^2^_3_ = 4.54, *P* = 0.209). Unsurprisingly, given the differences in the amount of pollen offered and stigma size by the flowers of each plant species, there were significant differences in pollen receipt among plant species (Figure 6b; GLMM for quantity: effect of plant species: χ^2^_3_ = 128.64, *P* < 0.0001; Figure S1b, GLMM for rate: effect of plant species: χ^2^_3_ = 66.75, *P* < 0.0001).

## DISCUSSION

Flower morphology that requires pollinators to learn how to extract rewards is often considered an adaptation that benefits the plant in terms of enhancing pollination (Darwin 1895; Laverty 1994; Lewis 1993; Jones and Agrawal 2017; Barker et al. 2018). We find that learning to manipulate flower morphology by pollen foraging bumble bees does not significantly affect benefits for the plant in terms of pollen deposition on flowers. For all four plant species tested, both the total amount and rate of pollen deposition did not differ between experienced and initially flower-naïve bumble bees. At the same time, we found strong benefits of learning for bees in terms of pollen collection. With experience, bees were much more likely to successfully extract pollen, collected 22% more pollen in their pollen baskets overall, and did so at a 24% greater rate. Furthermore, flower morphology significantly affected learning and memory, with some flower types being much more difficult to learn and remember than others. However, for no plant species did learning affect pollen deposition. Pollen foraging is often expected to have a greater (and negative) impact on pollination relative to nectar foraging (Wilson and Thomson 1991; Koski et al. 2018; Weinman et al. 2023). Assuming our results are broadly generalizable, difficult to learn flowers do not appear to be more costly to plants (Laverty 1980; Muth et al. 2015), at least in terms of reduced pollen transfer by inexperienced pollinators. Likewise, how learning affected pollen collection by bees did not differ among plant species with differing morphology, suggesting that difficult to learn flowers do not serve to protect floral rewards from being overexploited (Laverty 1994).

Why didn’t differences in learning and memory associated with different flower types affect pollen collection or pollen deposition? We assessed learning and memory of flower handling in multiple ways and not all were strongly affected by flower type. Indeed, our pollen foraging bees quickly reached an asymptote in their efficiency to handle any flower type (within 6-9 visits) and to locate the anthers after landing on any flower type (within 9-15 visits), on par with the fastest improvements observed for bees foraging for nectar on so-called morphologically ‘simple’ flowers (Laverty 1980, 1994, 1988). Although memory of flower handling efficiency by pollen foragers in this study was imperfect, relearning was likewise quick. Thus, modest effects of flower morphology on learning and memory of flower handling efficiency were likely a consequence of how few flower visits were required to learn and relearn efficient handling. Consequently, if flower handling efficiency drives patterns of reward collection and pollen transfer (as frequently suggested; e.g., Laverty 1980; Lewis 1993; Muth et al. 2015; Keaser et al. 2022), the lack of an effect of flower morphology is not surprising.

Given that cognitive constraints of efficiently handling flowers and locating the anthers were quite small, more slowly learned components of flower handling likely drove differences in pollen collection between experienced and inexperienced bees. Accordingly, flower handling motor routines and successful pollen extraction took many visits to learn and were strongly affected by flower type. Although flower handling motor routines have been described for inexperienced and experienced bees on different flower types (e.g., Laverty 1980; Chittka and Thomson 1997; Russell et al. 2018; Baek et al. 2023), the pace of learning handling techniques of different flower types has been unclear. Our results highlight that not all components of flower handling are learned at the same pace, with motor routines being learned particularly slowly. Additionally, substantial interindividual variation in the use of the modified motor routines suggests that experienced bees were still learning when trials were terminated. Assuming these bees were not close to proficiency, if we had allowed them many more flower visits to learn, we might have eventually observed effects of flower morphology on pollen collection and pollen transfer. Consistent with this hypothesis, Raine and Chittka (2007) found that bees were still improving their collection of pollen on a single plant species even after more than 300 flower visits. Altogether, determining conclusively how pollinator learning and memory affect pollen fate will require careful assessment of how different learned components of flower handling directly affect pollen collection and/or pollination.

Pollen collection by bees frequently reduces the amount of pollen available for pollination (Hargreaves et al. 2009; Russell et al. 2021; Weinman et al. 2023). Given that bees collected pollen more effectively and efficiently with experience, why didn’t learning simultaneously reduce pollen transfer? One possibility is that learning also made bees better at depositing pollen on flower stigmas, resulting in no overall change in pollen transfer for experienced versus inexperienced bees. For instance, modification of flower handling techniques with experience often involved the bee altering how it contacted the flower’s reproductive organs (Figure 5b), which could have resulted in more opportunities for pollen transfer to stigmas, even if less pollen overall was available for transfer to stigmas due to improved pollen collection. Similarly, learning might have reduced pollen wastage by the bee (i.e., pollen removed from the anthers, but not collected by the bee or deposited on stigmas; Parker et al. 2016; Minnaar et al. 2019). As a consequence, experienced bees would acquire more pollen on their bodies, which could explain increases in pollen collection without necessarily affecting pollen transfer. Quantifying how learning alters pollen wastage and contact with stigmas would thus be useful to more fully characterize effects of pollinator learning on pollination.

While we only quantified benefits to the plant in terms of pollen transfer, pollinator learning may have affected other components of pollination success. For instance, perhaps experienced bees depleted anthers more thoroughly, which would reduce pollen acquisition and transfer by subsequent visitors (Minnaar et al. 2019). A brief check of visited flowers in our trials revealed that they were not depleted of pollen, but we were unable to quantify whether flowers visited by experienced versus inexperienced bees were relatively more depleted. Additionally, we could not examine whether learning affected the quality of pollen transferred across visits, in terms of xenogamous versus same-flower self-pollen, which is likely affected by changes in flower handling time and the sequence of flower visits (Robertson 1992; Karron et al. 2009; Horsburg et al. 2011; Minnaar et al. 2019; Minnaar and Anderson 2018). Finally, effects of learning on pollen deposition may not have been observed if bees delivered much more pollen to stigmas than these reproductive structures could physically receive. However, this possibility is unlikely: for each plant species, the maximum quantity of pollen on stigmas after behavioral trials was much greater than the average quantity of pollen transferred to stigmas (greater by 2.1x, 2.7x, 3.4x, and 4.7x for *E. affine*, *P. campanularia*, *E. guttata,* and *S. elaeagnifolium* respectively), suggesting that on average, stigmas were not saturated with pollen.

In conclusion, flower morphologies that conceal pollen rewards to a greater degree are overall more difficult to learn and costs of learning flower handling depend on flower type, just as has been found in a nectar reward context (Laverty 1980, 1994; Gegear and Laverty 1995; Muth et al. 2015; Keaser et al. 2022). Additionally, we present rare quantitative evidence of how pollinator learning influences pollen fate, demonstrating strong positive effects for reward collection, but no effects on pollen transfer to flowers. Our results thus confirm and extend those of Ramos et al. (2017), who also found no effect of flower handling experience by nectar-foraging butterflies on pollinia transfer. Finally, our work has particular relevance for understanding why pollinators such as bees often exhibit floral fidelity (i.e., flower constancy), a pattern of behavior hypothesized to benefit pollinators and to drive flower evolution (Waser 1986; Chittka et al. 1999; Gruter and Ratnieks 2011; Muth et al. 2015; Ramos et al. 2017; Papaj and Russell 2024). Costs associated with learning a given flower type are often thought to facilitate floral fidelity, but time penalties incurred while learning to efficiently handle flowers are often small (as we found here) and thus likely insufficient to facilitate floral fidelity (Woodward and Laverty 1992; Gegear and Laverty 1995; Chittka et al. 1999). Although costs of learning motor routines in terms of reward collection have yet to be examined in the context of floral fidelity (but see Chittka and Thompson 1997), our results suggest that these costs are far larger and thus have the potential to facilitate floral fidelity.

## Supporting information

Data and R script used in analyses

## ACKNOWLEDGEMENTS

We are grateful to Koppert Biologicals for bee colonies, Abilene Mosher for greenhouse care, and Russell lab members for discussion. We acknowledge this work was performed on unceded traditional territory of the Kiikaapoi, Sioux, and Osage.

## DECLARATIONS

### ETHICS APPROVAL

All bumble bee experimentation was carried out in accordance with the legal and ethical standards of the USA.

### CONSENT FOR PUBLICATION

Not applicable

### DATA AVAILABILITY STATEMENT

The datasets supporting this article are available as electronic supplementary material.

## SUPPLEMENTARY

**Figure S1.**
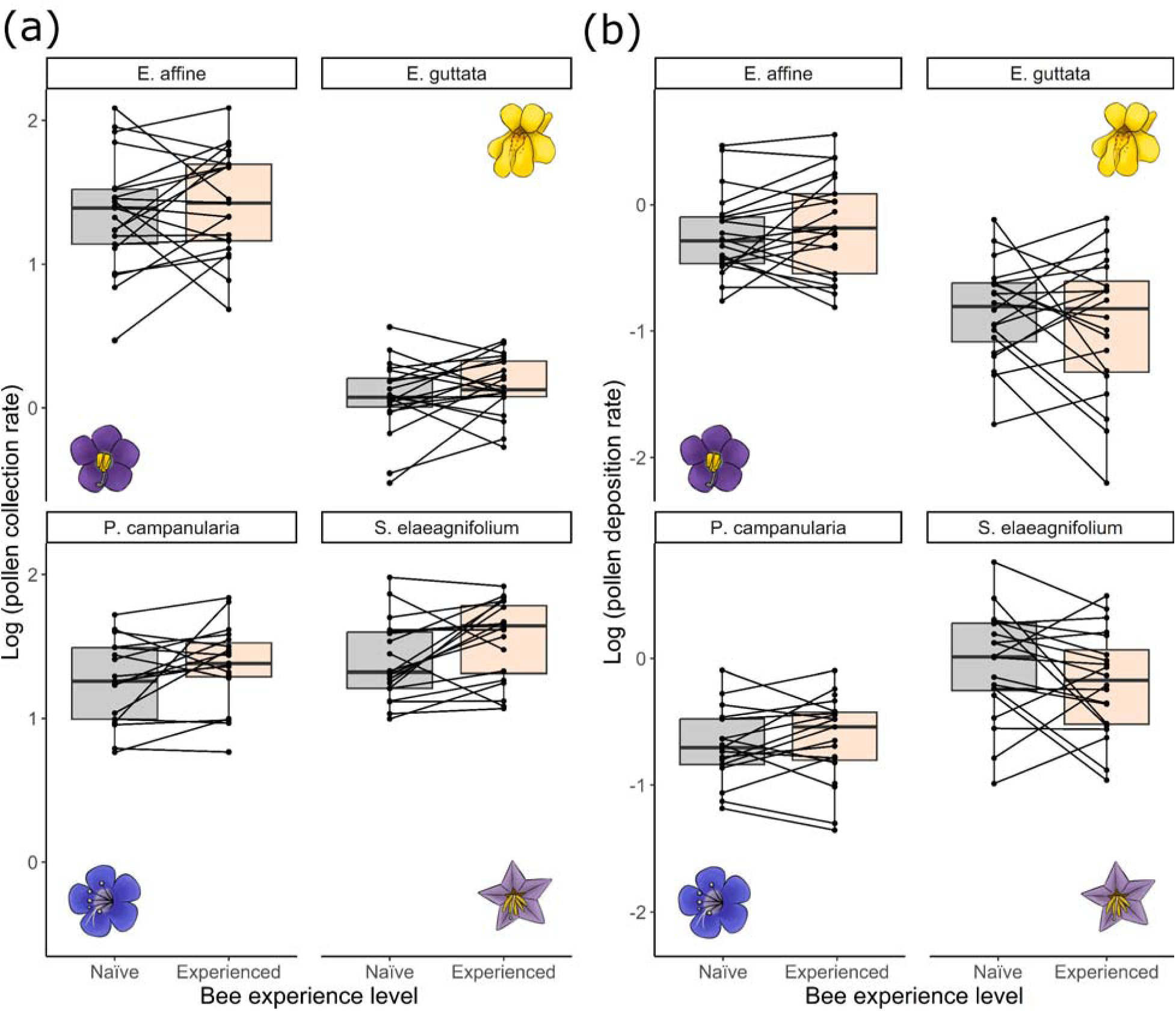
The rate of pollen (a) collected into the pollen baskets (corbiculae) or (b) transferred to flower stigmas, as influenced by bee experience and plant species. Plotted as boxplots with raw data indicated via dots joined by lines showing how the amount of pollen collected or transferred by individual bees changed with experience. *N* = 21, 20, 17, 20 initially naïve bees and *N* = 21, 20, 19, 20 experienced bees foraging on *E. affine*, *E. guttata, P. campanularia*, and *S. elaeagnifolium*, respectively.

## Notes

### Competing Interest Statement

The authors have declared no competing interest.

