## Supplementary material for "Learning to handle flowers increases pollen collection benefits for bees but does not affect pollination success for plants": Data and R script used in analyses: Analysis_learning-affects-pollination-benefits_5-6-2024_final.R

```
# Created on 7-3-2022
# Updated data on 7-9-2022 to include motor routines (with errors)
# Updated with all but two EXAF bees 7-16-2022
# Everything is updated 2023
# Updated with new graphs 4-28-2023
# Updated with failure data 5-22-2023
# Updated with handling time traces 5-24-2023
# Updated with pollen deposited/collected per second
# Changed name from M. guttatus to E. guttata


#Name of data file:
FILENAME1 <- "AnalysisData_All-Trials-Summary-Data_20_5-28-2023.csv"
FILENAME2 <- "AnalysisData_Handling-Time-Traces-PHCA_5-24-2023.csv" # For graphs only
FILENAME3 <- "AnalysisData_Handling-Time-Traces-EXAF_5-24-2023.csv" # For graphs only
FILENAME4 <- "AnalysisData_Handling-Time-Traces-SOEL_5-24-2023.csv" # For graphs only
FILENAME5 <- "AnalysisData_Handling-Time-Traces-MIGU_5-24-2023.csv" # For graphs only
FILENAME6 <- "AnalysisData_Naive-Bees_Efficiency_5-28-2023.csv"


#Sets working directory automatically to where R script is saved
setwd(dirname(rstudioapi::getActiveDocumentContext()$path))

# Set the directory your plots will be in:
PLOTDIR <- dirname(rstudioapi::getActiveDocumentContext()$path)

#Load data:
setwd(dirname(rstudioapi::getActiveDocumentContext()$path)); dataset1 <- read.table(FILENAME1, sep=",", header=TRUE)
setwd(dirname(rstudioapi::getActiveDocumentContext()$path)); dataset2 <- read.table(FILENAME2, sep=",", header=TRUE)
setwd(dirname(rstudioapi::getActiveDocumentContext()$path)); dataset3 <- read.table(FILENAME3, sep=",", header=TRUE)
setwd(dirname(rstudioapi::getActiveDocumentContext()$path)); dataset4 <- read.table(FILENAME4, sep=",", header=TRUE)
setwd(dirname(rstudioapi::getActiveDocumentContext()$path)); dataset5 <- read.table(FILENAME5, sep=",", header=TRUE)
setwd(dirname(rstudioapi::getActiveDocumentContext()$path)); dataset6 <- read.table(FILENAME6, sep=",", header=TRUE)


#libraries
library("mgcv")
library("lmerTest")
library("mclust")
library("DescTools")
library("lme4")
library("lsmeans")
library("multcomp")
library("DHARMa")
library("car")
library("glmmTMB")
library("lme4")
library("effects") #For plots
library("ggplot2") #For plots
library("RVAideMemoire")
library("performance") #install.packages("performance") #To use check_model assumptions
library("see") #install.packages("see") #To use check_model assumptions
library("gridExtra") #install.packages("gridExtra") #To use check_model assumptions
library("ARTool")
library("sjPlot")

R.Version()
citation("R Package Name Here") 


#### Efficiency of handling time, relative to experienced bees ####
# Naive bees

# Run the model
kruskal.test(NumVisits_HandTimeEfficient ~ Species, data = dataset6)

# Kruskal-Wallis chi-squared = 3.9869, df = 3, p-value = 0.2629

# Run the model
kruskal.test(Efficiency_FirstVisitHandTime ~ Species, data = dataset6)

# Kruskal-Wallis chi-squared = 14.535, df = 3, p-value = 0.00226

# Pairwise comparisons with corrections for multiple testing
pairwise.wilcox.test(dataset6$Efficiency_FirstVisitHandTime, dataset6$Species, p.adjust.method = "BH")
#       EXAF  MIGU  PHCA 
# MIGU 0.015 -     -    
# PHCA 0.015 0.964 -    
# SOEL 0.653 0.015 0.015

# Efficiency
#     MIGU a     EXAF b       SOEL b     PHCA a
# 0.411955835	0.739807165	1.012586526	0.395547202


# Run the model
kruskal.test(Slope_EfficientHandTime ~ Species, data = dataset6)

# Kruskal-Wallis chi-squared = 0.33911, df = 3, p-value = 0.9525


# Examine whether slope and efficiency are correlated
# Run the model
Slope.vs.Efficiency.HT <- lm(Efficiency_FirstVisitHandTime ~ Slope_EfficientHandTime, data = dataset6)

# Examine the model
summary(Slope.vs.Efficiency.HT)
Anova(Slope.vs.Efficiency.HT)

#                           Sum Sq Df F value Pr(>F)
# Slope_EfficientHandTime  0.0231  1  0.0933 0.7609


#### Efficiency of latency to begin extracting pollen from anthers, relative to experienced bees ####
# Naive bees

# Run the model
kruskal.test(NumVisits_AnthLatEfficient ~ Species, data = dataset6)

# Kruskal-Wallis chi-squared = 0.026965, df = 3, p-value = 0.9988

# Run the model
kruskal.test(Efficiency_FirstVisitAnthLate ~ Species, data = dataset6)

# Kruskal-Wallis chi-squared = 4.0739, df = 3, p-value = 0.2536

# Run the model
kruskal.test(Slope_EfficientAnthLate ~ Species, data = dataset6)

# Kruskal-Wallis chi-squared = 5.7079, df = 3, p-value = 0.1267


# Examine whether slope and efficiency are correlated
# Run the model
Slope.vs.Efficiency.AL <- lm(Efficiency_FirstVisitAnthLate ~ Slope_EfficientAnthLate, data = dataset6)

# Examine the model
summary(Slope.vs.Efficiency.AL)
Anova(Slope.vs.Efficiency.AL)

# Slope_EfficientAnthLate  0.7168  1  2.6539 0.1083


# Question
#### Is learning to handle flowers (time in seconds) affected by plant species and treatment? ####
# Parameters 'a' and 'b' from Wright's cumulative learning model on first 20 visits


# Here we examine how 'a' differs for first or second day only

# Run the model
LearnSlope.DayOne <- aov(LearnSlopeDayOne ~ SpeciesDayOne, data = dataset1)
LearnSlope.DayTwo <- aov(LearnSlopeDayTwo ~ SpeciesDayTwo, data = dataset1)

# Examine the model
summary(LearnSlope.DayOne)

#               Df   Sum Sq   Mean Sq F value   Pr(>F)    
# SpeciesDayOne  3 0.003544 0.0011815   7.047 0.000313 ***

summary(LearnSlope.DayTwo)

#               Df   Sum Sq   Mean Sq F value Pr(>F)
# SpeciesDayTwo  3 0.000668 0.0002226   1.161   0.33

# Post-hoc test
emmeans(LearnSlope.DayOne, list(pairwise ~ SpeciesDayOne), adjust = "tukey")

# SpeciesDayOne   emmean      SE df lower.CL  upper.CL
# EXAF          -0.01200 0.00283 74  -0.0176 -0.006365
# MIGU          -0.02438 0.00290 74  -0.0301 -0.018611
# PHCA          -0.01759 0.00314 74  -0.0239 -0.011337
# SOEL          -0.00639 0.00290 74  -0.0122 -0.000626

# $`pairwise differences of SpeciesDayOne`
# 1           estimate      SE df t.ratio p.value
# EXAF - MIGU  0.01238 0.00405 74   3.061  0.0159
# EXAF - PHCA  0.00560 0.00422 74   1.325  0.5499
# EXAF - SOEL -0.00560 0.00405 74  -1.384  0.5131
# MIGU - PHCA -0.00679 0.00427 74  -1.589  0.3913
# MIGU - SOEL -0.01799 0.00409 74  -4.392  0.0002
# PHCA - SOEL -0.01120 0.00427 74  -2.622  0.0508

emmeans(LearnSlope.DayTwo, list(pairwise ~ SpeciesDayTwo), adjust = "tukey")

#  SpeciesDayTwo   emmean      SE df lower.CL  upper.CL
# EXAF          -0.00556 0.00302 75  -0.0116  0.000457    No learning (encompasses zero)
# MIGU          -0.00832 0.00318 75  -0.0146 -0.001993
# PHCA          -0.01350 0.00318 75  -0.0198 -0.007172
# SOEL          -0.00768 0.00310 75  -0.0139 -0.001517

# $`pairwise differences of SpeciesDayTwo`
# 1            estimate      SE df t.ratio p.value
# EXAF - MIGU  0.002759 0.00438 75   0.629  0.9223
# EXAF - PHCA  0.007938 0.00438 75   1.811  0.2765
# EXAF - SOEL  0.002123 0.00433 75   0.491  0.9609
# MIGU - PHCA  0.005179 0.00449 75   1.153  0.6582
# MIGU - SOEL -0.000636 0.00444 75  -0.143  0.9989
# PHCA - SOEL -0.005815 0.00444 75  -1.311  0.5590

# Here we examine how 'b' differs for first or second day only

# Run the model
LearnIntercept.DayOne <- aov(sqrt(LearnInterceptDayOne) ~ SpeciesDayOne, data = dataset1)
LearnIntercept.DayTwo <- aov(sqrt(LearnInterceptDayTwo) ~ SpeciesDayTwo, data = dataset1)

# Examine the model
summary(LearnIntercept.DayOne)

# SpeciesDayOne  3  201.8   67.28   30.02 8.35e-13 ***

summary(LearnIntercept.DayTwo)

# SpeciesDayTwo  3  63.08  21.026   10.47 8.01e-06 ***

# Post-hoc test
emmeans(LearnIntercept.DayOne, list(pairwise ~ SpeciesDayOne), adjust = "tukey")

#  SpeciesDayOne emmean    SE df lower.CL upper.CL
# EXAF            4.78 0.327 74     4.13     5.44
# MIGU            7.47 0.335 74     6.80     8.14
# PHCA            3.26 0.363 74     2.54     3.99
# SOEL            3.81 0.335 74     3.14     4.47

# $`pairwise differences of SpeciesDayOne`
# 1           estimate    SE df t.ratio p.value
# EXAF - MIGU   -2.686 0.468 74  -5.743  <.0001
# EXAF - PHCA    1.520 0.488 74   3.112  0.0138
# EXAF - SOEL    0.978 0.468 74   2.091  0.1656
# MIGU - PHCA    4.206 0.494 74   8.517  <.0001
# MIGU - SOEL    3.664 0.473 74   7.740  <.0001
# PHCA - SOEL   -0.542 0.494 74  -1.097  0.6922

emmeans(LearnIntercept.DayTwo, list(pairwise ~ SpeciesDayTwo), adjust = "tukey")

# SpeciesDayTwo emmean    SE df lower.CL upper.CL
# EXAF            4.11 0.309 74     3.49     4.72
# MIGU            5.14 0.334 74     4.47     5.80
# PHCA            2.55 0.325 74     1.90     3.20
# SOEL            4.01 0.317 74     3.38     4.65

# $`pairwise differences of SpeciesDayTwo`
# 1           estimate    SE df t.ratio p.value
# EXAF - MIGU  -1.0288 0.455 74  -2.261  0.1168
# EXAF - PHCA   1.5557 0.449 74   3.468  0.0048
# EXAF - SOEL   0.0917 0.443 74   0.207  0.9968
# MIGU - PHCA   2.5845 0.466 74   5.546  <.0001
# MIGU - SOEL   1.1205 0.460 74   2.434  0.0795
# PHCA - SOEL  -1.4640 0.454 74  -3.226  0.0099


#### Here we examine how 'a' or 'b' differ from each other with experience ####

# Run the model
Learn.Slope <- glmmTMB((LearnSlope) ~ Treatment * Species + (1|BeeID), data = dataset1) # Would not coverge with ColonyID
Learn.Intercept <- glmmTMB(sqrt(LearnIntercept) ~ Treatment * Species + (1|ColonyID/BeeID), data = dataset1)

# Examine the model
summary(Learn.Slope)

summary(Learn.Intercept)


# Check Type II sums of squares, for Wald Tests
Anova(Learn.Slope)

Anova(Learn.Intercept)

#Post-hoc test to see significance of the pairs (Treatment 1 vs Treatment 2)
#summary(glht(Learn.Slope, linfct=mcp(Species="Tukey"))) #Use lmer in the model above to make this work

# Response: (LearnSlope)
#                     Chisq Df Pr(>Chisq)  
# Treatment         15.166  1  9.847e-05 ***
# Species           11.706  3   0.008462 ** 
# Treatment:Species 14.501  3   0.002297 ** 

# Response: sqrt(LearnIntercept)
#                    Chisq Df Pr(>Chisq)    
# Treatment         28.176  1  1.108e-07 ***
# Species           80.024  3  < 2.2e-16 ***
# Treatment:Species 30.262  3  1.216e-06 ***

# Post-hoc test
emmeans(Learn.Intercept, list(pairwise ~ Species * Treatment), adjust = "tukey")

# Sjplot effect sizes, data, and predicted values
plot_model(Learn.Intercept)
plot_model(Learn.Intercept, type="pred", terms=c("Treatment"), show.data = T) #
plot_model(Learn.Intercept, type="pred", terms=c("Species", "Treatment"), show.data = T) #

plot_model(Learn.Slope)
plot_model(Learn.Slope, type="pred", terms=c("Treatment"), show.data = T) #
plot_model(Learn.Slope, type="pred", terms=c("Species", "Treatment"), show.data = T) #


#Analysis of Deviance Table (Type II Wald chisquare tests)

# Response: (LearnSlope)
#             Chisq Df Pr(>Chisq)
# Treatment 1.6776  1     0.1952
# Species   1.4074  2     0.4948

#                      Chisq Df Pr(>Chisq)    
# Treatment          6.2861  1    0.01217 *  
# Species           30.9256  2  1.926e-07 ***
# Treatment:Species  5.3156  2    0.07010

# Model assumptions via DHARMa
simulationOutput1 <- simulateResiduals(fittedModel = Learn.Intercept, n = 10000) #Learn.Slope
plot(simulationOutput1) #QQ plots good after remove O8_Two outlier for slope, W8 / W11_Two for intercept
testUniformity(simulationOutput = simulationOutput1) #Uniformity good 
testDispersion(simulationOutput1) #Dispersion good
testZeroInflation(simulationOutput1) #Zero inflation good

# Figure with paired data

# Rename the figure legends
dataset1$Treatment = plyr::mapvalues(dataset1$Treatment, from = c("DayOne", "DayTwo"), to = c("Naïve", "Experienced"))
dataset1$Species = plyr::mapvalues(dataset1$Species, from = c("EXAF", "MIGU", "PHCA", "SOEL"), 
                                   to = c("E. affine", "E. guttata", "P. campanularia", "S. elaeagnifolium"))

# Reorder the experience levels (treatment variables)
dataset1$Treatment <- factor(dataset1$Treatment, levels = c("Naïve","Experienced"))

# Plot figure
Figure2a <- ggplot(dataset1, aes(x = Treatment, y = LearnSlope)) + 
  geom_boxplot(aes(fill = Treatment), alpha = .2) +
  geom_line(aes(group = BeeID)) + 
  geom_point(size = 1) + 
  facet_wrap(~ Species) +
  ylab("Slope of the learning curve") + xlab("Bee experience level") + theme_classic() + 
  theme(text=element_text(size=14)) +
  scale_color_viridis_d(name = "Experience \nlevel", end = 0.7, option = "inferno") + 
  scale_fill_viridis_d(name = "Experience \nlevel", end =0.7, option = 'inferno') +
  theme(legend.position  = c(-1.0, -1.0), legend.direction = "horizontal", 
        legend.background = element_rect(fill="transparent", colour = NA),
        legend.title = element_text(size = 0), legend.text = element_text(size = 0),
        panel.spacing.x = unit(1, "lines"), 
        plot.margin = margin(c(0,0, 0, 0), unit = "cm"))
plot(Figure2a) #To see the figure in R

# Set date
DATE <- "4-28-2023"

tiff(filename=paste("Fig2a_Learn-Slope_", DATE, ".tif", sep=""), units="mm", width=120, height=200, res=1200)

plot(Figure2a) #To export it

setwd(PLOTDIR); 

dev.off()


# Rename the figure legends
dataset1$Treatment = plyr::mapvalues(dataset1$Treatment, from = c("DayOne", "DayTwo"), to = c("Naïve", "Experienced"))
dataset1$Species = plyr::mapvalues(dataset1$Species, from = c("EXAF", "MIGU", "PHCA", "SOEL"), 
                                   to = c("E. affine", "E. guttata", "P. campanularia", "S. elaeagnifolium"))

# Reorder the experience levels (treatment variables)
dataset1$Treatment <- factor(dataset1$Treatment, levels = c("Naïve","Experienced"))

# Plot figure
Figure2b <- ggplot(dataset1, aes(x = Treatment, y = sqrt(LearnIntercept))) + 
  geom_boxplot(aes(fill = Treatment), alpha = .2) +
  geom_line(aes(group = BeeID)) + 
  geom_point(size = 1) + 
  facet_wrap(~ Species) +
  ylab("Sqrt(intercept of the learning curve)") + xlab("Bee experience level") + theme_classic() + 
  theme(text=element_text(size=14)) +
  scale_color_viridis_d(name = "Experience \nlevel", end = 0.7, option = "inferno") + 
  scale_fill_viridis_d(name = "Experience \nlevel", end =0.7, option = 'inferno') +
  theme(legend.position  = c(-1.0, -1.0), legend.direction = "horizontal", 
        legend.background = element_rect(fill="transparent", colour = NA),
        legend.title = element_text(size = 0), legend.text = element_text(size = 0),
        panel.spacing.x = unit(1, "lines"), 
        plot.margin = margin(c(0,0, 0, 0), unit = "cm"))
plot(Figure2b) #To see the figure in R

# Set date
DATE <- "4-28-2023"

tiff(filename=paste("Fig2b_Learn-Intercept_", DATE, ".tif", sep=""), units="mm", width=120, height=200, res=1200)

plot(Figure2b) #To export it

setwd(PLOTDIR); 

dev.off()


#_______________________________________________________________________________________________________________________________#


# Question
#### Is learning to find the anthers (time in seconds) affected by plant species and treatment? ####
# Parameters 'a' and 'b' from Wright's cumulative learning model on first 20 visits


# Here we examine how 'a' differs for first or second day only

# Run the model
Anth.LearnSlope.DayOne <- aov(AnthLearnSlopeDayOne ~ SpeciesDayOne, data = dataset1)
Anth.LearnSlope.DayTwo <- aov(AnthLearnSlopeDayTwo ~ SpeciesDayTwo, data = dataset1)

# Examine the model
summary(Anth.LearnSlope.DayOne)

#               Df   Sum Sq   Mean Sq F value   Pr(>F)    
# SpeciesDayOne  3 0.002542 0.0008473   3.664 0.0161 *

summary(Anth.LearnSlope.DayTwo)

#               Df   Sum Sq   Mean Sq F value Pr(>F)
# SpeciesDayTwo  3 0.001439 0.0004795   3.606 0.0172 *

# Post-hoc test
emmeans(Anth.LearnSlope.DayOne, list(pairwise ~ SpeciesDayOne), adjust = "tukey")

# SpeciesDayOne   emmean      SE df lower.CL  upper.CL
# EXAF          -0.00824 0.00332 74  -0.0149 -0.001630
# MIGU          -0.01948 0.00340 74  -0.0263 -0.012704
# PHCA          -0.00423 0.00369 74  -0.0116  0.003120    No learning (encompasses zero)
# SOEL          -0.00761 0.00340 74  -0.0144 -0.000834

# $`pairwise differences of SpeciesDayOne`
# 1           estimate      SE df t.ratio p.value
# EXAF - MIGU  0.011237 0.00475 74   2.365  0.0930
# EXAF - PHCA -0.004013 0.00496 74  -0.809  0.8501
# EXAF - SOEL -0.000633 0.00475 74  -0.133  0.9991
# MIGU - PHCA -0.015251 0.00502 74  -3.040  0.0169
# MIGU - SOEL -0.011870 0.00481 74  -2.468  0.0735
# PHCA - SOEL  0.003381 0.00502 74   0.674  0.9067

emmeans(Anth.LearnSlope.DayTwo, list(pairwise ~ SpeciesDayTwo), adjust = "tukey")

#  SpeciesDayTwo   emmean      SE df lower.CL  upper.CL
# EXAF          -0.00571 0.00252 75 -0.01073 -0.000701
# MIGU          -0.00780 0.00265 75 -0.01308 -0.002535
# PHCA          -0.00314 0.00265 75 -0.00841  0.002133    No learning (encompasses zero)
# SOEL          -0.01464 0.00258 75 -0.01977 -0.009498

# $`pairwise differences of SpeciesDayTwo`
# 1            estimate      SE df t.ratio p.value
# EXAF - MIGU  0.00209 0.00365 75   0.573  0.9400
# EXAF - PHCA -0.00258 0.00365 75  -0.706  0.8945
# EXAF - SOEL  0.00892 0.00360 75   2.476  0.0720
# MIGU - PHCA -0.00467 0.00374 75  -1.248  0.5988
# MIGU - SOEL  0.00683 0.00369 75   1.849  0.2591
# PHCA - SOEL  0.01150 0.00369 75   3.112  0.013

# Here we examine how 'b' differs for first or second day only

# Run the model
Anth.LearnIntercept.DayOne <- aov(sqrt(AnthLearnInterceptDayOne) ~ SpeciesDayOne, data = dataset1)
Anth.LearnIntercept.DayTwo <- aov(sqrt(AnthLearnInterceptDayTwo) ~ SpeciesDayTwo, data = dataset1)

# Examine the model
summary(Anth.LearnIntercept.DayOne)

# SpeciesDayOne  3  43.24  14.413   36.71 1.22e-14 ***

summary(Anth.LearnIntercept.DayTwo)

# SpeciesDayTwo  3  12.03   4.008   15.93 3.96e-08 ***

# Post-hoc test
emmeans(Anth.LearnIntercept.DayOne, list(pairwise ~ SpeciesDayOne), adjust = "tukey")

#  SpeciesDayOne emmean    SE df lower.CL upper.CL
# EXAF            1.73 0.137 74     1.46     2.01
# MIGU            3.34 0.140 74     3.06     3.62
# PHCA            1.43 0.152 74     1.12     1.73
# SOEL            1.78 0.140 74     1.50     2.06

# $`pairwise differences of SpeciesDayOne`
# 1           estimate    SE df t.ratio p.value
# EXAF - MIGU  -1.6051 0.196 74  -8.198  <.0001
# EXAF - PHCA   0.3082 0.204 74   1.507  0.4383
# EXAF - SOEL  -0.0499 0.196 74  -0.255  0.9942
# MIGU - PHCA   1.9133 0.207 74   9.256  <.0001
# MIGU - SOEL   1.5552 0.198 74   7.849  <.0001
# PHCA - SOEL  -0.3581 0.207 74  -1.732  0.3146

emmeans(Anth.LearnIntercept.DayTwo, list(pairwise ~ SpeciesDayTwo), adjust = "tukey")

# SpeciesDayTwo emmean    SE df lower.CL upper.CL
# EXAF            1.60 0.109 76     1.38     1.82
# MIGU            2.45 0.112 76     2.22     2.67
# PHCA            1.42 0.115 76     1.19     1.65
# SOEL            1.78 0.112 76     1.56     2.01

# $`pairwise differences of SpeciesDayTwo`
# 1           estimate    SE df t.ratio p.value
# EXAF - MIGU   -0.850 0.157 76  -5.426  <.0001
# EXAF - PHCA    0.180 0.159 76   1.134  0.6701
# EXAF - SOEL   -0.187 0.157 76  -1.194  0.6324
# MIGU - PHCA    1.030 0.161 76   6.412  <.0001
# MIGU - SOEL    0.663 0.159 76   4.181  0.0004
# PHCA - SOEL   -0.367 0.161 76  -2.285  0.1106


#### Here we examine how 'a' or 'b' differ from each other with experience ####

# Run the model
Anth.Learn.Slope <- glmmTMB((AntherLearnSlope) ~ Treatment * Species + (1|BeeID), data = dataset1) # Would not coverge with ColonyID
Anth.Learn.Intercept <- glmmTMB(sqrt(AntherLearnIntercept) ~ Treatment * Species + (1|ColonyID/BeeID), data = dataset1)

# Examine the model
summary(Anth.Learn.Slope)

summary(Anth.Learn.Intercept)


# Check Type II sums of squares, for Wald Tests
Anova(Anth.Learn.Slope)

Anova(Anth.Learn.Intercept)

#Post-hoc test to see significance of the pairs (Treatment 1 vs Treatment 2)
#summary(glht(Learn.Slope, linfct=mcp(Species="Tukey"))) #Use lmer in the model above to make this work

# Response: (AntherLearnSlope)
#                     Chisq Df Pr(>Chisq)  
# Treatment          1.2027  1   0.272788   
# Species           10.7485  3   0.013167 * 
# Treatment:Species 12.8497  3   0.004973 **

# Response: sqrt(AntherLearnIntercept)
#                    Chisq Df Pr(>Chisq)    
# Treatment          10.809  1  0.0010102 ** 
# Species           140.703  3  < 2.2e-16 ***
# Treatment:Species  21.049  3  0.0001029 ***

# Post-hoc test
emmeans(Anth.Learn.Intercept, list(pairwise ~ Treatment * Species), adjust = "tukey")

# Sjplot effect sizes, data, and predicted values
plot_model(Anth.Learn.Intercept)
plot_model(Anth.Learn.Intercept, type="pred", terms=c("Treatment"), show.data = T) #
plot_model(Anth.Learn.Intercept, type="pred", terms=c("Species", "Treatment"), show.data = T) #

plot_model(Anth.Learn.Slope)
plot_model(Anth.Learn.Slope, type="pred", terms=c("Treatment"), show.data = T) #
plot_model(Anth.Learn.Slope, type="pred", terms=c("Species", "Treatment"), show.data = T) #


#Analysis of Deviance Table (Type II Wald chisquare tests)

# Response: (LearnSlope)
#             Chisq Df Pr(>Chisq)
# Treatment 1.6776  1     0.1952
# Species   1.4074  2     0.4948

#                      Chisq Df Pr(>Chisq)    
# Treatment          6.2861  1    0.01217 *  
# Species           30.9256  2  1.926e-07 ***
# Treatment:Species  5.3156  2    0.07010

# Model assumptions via DHARMa
simulationOutput1 <- simulateResiduals(fittedModel = Anth.Learn.Slope, n = 10000) #Learn.Slope
plot(simulationOutput1) #QQ plots good after remove O8_Two outlier for slope
testUniformity(simulationOutput = simulationOutput1) #Uniformity good 
testDispersion(simulationOutput1) #Dispersion good
testZeroInflation(simulationOutput1) #Zero inflation good

# Figure with paired data

# Rename the figure legends
dataset1$Treatment = plyr::mapvalues(dataset1$Treatment, from = c("DayOne", "DayTwo"), to = c("Naïve", "Experienced"))
dataset1$Species = plyr::mapvalues(dataset1$Species, from = c("EXAF", "MIGU", "PHCA", "SOEL"), 
                                   to = c("E. affine", "E. guttata", "P. campanularia", "S. elaeagnifolium"))

# Reorder the experience levels (treatment variables)
dataset1$Treatment <- factor(dataset1$Treatment, levels = c("Naïve","Experienced"))

# Plot figure
Figure3a <- ggplot(dataset1, aes(x = Treatment, y = AntherLearnSlope)) + 
  geom_boxplot(aes(fill = Treatment), alpha = .2) +
  geom_line(aes(group = BeeID)) + 
  geom_point(size = 1) + 
  facet_wrap(~ Species) +
  ylab("Slope of the learning curve") + xlab("Bee experience level") + theme_classic() + 
  theme(text=element_text(size=14)) +
  scale_color_viridis_d(name = "Experience \nlevel", end = 0.7, option = "inferno") + 
  scale_fill_viridis_d(name = "Experience \nlevel", end =0.7, option = 'inferno') +
  theme(legend.position  = c(-1.0, -1.0), legend.direction = "horizontal", 
        legend.background = element_rect(fill="transparent", colour = NA),
        legend.title = element_text(size = 0), legend.text = element_text(size = 0),
        panel.spacing.x = unit(1, "lines"), 
        plot.margin = margin(c(0,0, 0, 0), unit = "cm"))
plot(Figure3a) #To see the figure in R

# Set date
DATE <- "5-4-2023"

tiff(filename=paste("Fig3a_Anth-Learn-Slope_", DATE, ".tif", sep=""), units="mm", width=120, height=200, res=1200)

plot(Figure3a) #To export it

setwd(PLOTDIR); 

dev.off()


# Rename the figure legends
dataset1$Treatment = plyr::mapvalues(dataset1$Treatment, from = c("DayOne", "DayTwo"), to = c("Naïve", "Experienced"))
dataset1$Species = plyr::mapvalues(dataset1$Species, from = c("EXAF", "MIGU", "PHCA", "SOEL"), 
                                   to = c("E. affine", "E. guttata", "P. campanularia", "S. elaeagnifolium"))

# Reorder the experience levels (treatment variables)
dataset1$Treatment <- factor(dataset1$Treatment, levels = c("Naïve","Experienced"))

# Plot figure
Figure3b <- ggplot(dataset1, aes(x = Treatment, y = sqrt(AntherLearnIntercept))) + 
  geom_boxplot(aes(fill = Treatment), alpha = .2) +
  geom_line(aes(group = BeeID)) + 
  geom_point(size = 1) + 
  facet_wrap(~ Species) +
  ylab("Sqrt (intercept of the learning curve)") + xlab("Bee experience level") + theme_classic() + 
  theme(text=element_text(size=14)) +
  scale_color_viridis_d(name = "Experience \nlevel", end = 0.7, option = "inferno") + 
  scale_fill_viridis_d(name = "Experience \nlevel", end =0.7, option = 'inferno') +
  theme(legend.position  = c(-1.0, -1.0), legend.direction = "horizontal", 
        legend.background = element_rect(fill="transparent", colour = NA),
        legend.title = element_text(size = 0), legend.text = element_text(size = 0),
        panel.spacing.x = unit(1, "lines"), 
        plot.margin = margin(c(0,0, 0, 0), unit = "cm"))
plot(Figure3b) #To see the figure in R

# Set date
DATE <- "5-4-2023"

tiff(filename=paste("Fig3b_Anth-Learn-Intercept_", DATE, ".tif", sep=""), units="mm", width=120, height=200, res=1200)

plot(Figure3b) #To export it

setwd(PLOTDIR); 

dev.off()


#_______________________________________________________________________________________________________________________________#


# Question
#### Is the proportion of time spent on novel handling behaviors over all visits affected by treatment? ####
# Could not do it per visit and analyze learning rate, since no models would converge
# Avg of first 20 visits analyzed; to match with Wright's learning curve data

# Run the model
# Nested random effects; Bee within Colony
Novel.Motor.Routine <- glmmTMB(sqrt(MeanPropMotRoutine*100+1) ~ Species * Treatment  + (1|ColonyID/BeeID), data = dataset1)

# Examine the model
summary(Novel.Motor.Routine)

# Check Type II sums of squares, for Wald Tests
Anova(Novel.Motor.Routine)

# Response: PropVisitUsingBehav
#                     Chisq Df Pr(>Chisq)    
# Species           99.6423  2  < 2.2e-16 ***
# Treatment         24.8238  1  6.282e-07 ***
# Species:Treatment  2.2166  2     0.3301

emmeans(Novel.Motor.Routine, list(pairwise ~ Species * Treatment), adjust = "tukey")

# Sjplot effect sizes, data, and predicted values
plot_model(Novel.Motor.Routine)
plot_model(Novel.Motor.Routine, type="pred", terms=c("Treatment"), show.data = T) #
plot_model(Novel.Motor.Routine, type="pred", terms=c("Species", "Treatment"), show.data = T) #

# Model assumptions via DHARMa
simulationOutput1 <- simulateResiduals(fittedModel = Novel.Motor.Routine, n = 10000)
plot(simulationOutput1) #QQ plot is good
testUniformity(simulationOutput = simulationOutput1) #Uniformity good 
testDispersion(simulationOutput1) #Dispersion good
testZeroInflation(simulationOutput1) #Zero inflation good


# Figure with paired data

# Rename the figure legends
dataset1$Treatment = plyr::mapvalues(dataset1$Treatment, from = c("DayOne", "DayTwo"), to = c("Naïve", "Experienced"))
dataset1$Species = plyr::mapvalues(dataset1$Species, from = c("EXAF", "MIGU", "SOEL"), 
                                   to = c("E. affine", "E. guttata", "S. elaeagnifolium"))

# Reorder the experience levels (treatment variables)
dataset1$Treatment <- factor(dataset1$Treatment, levels = c("Naïve","Experienced"))

# Plot figure
Figure4a <- ggplot(dataset1, aes(x = Treatment, y = sqrt(MeanPropMotRoutine*100+1))) + 
  geom_boxplot(aes(fill = Treatment), alpha = .2) +
  geom_line(aes(group = BeeID)) + 
  geom_point(size = 1) + 
  facet_wrap(~ Species) +
  ylab("Sqrt (percent use of modified handling techniques)") + xlab("Bee experience level") + theme_classic() + 
  theme(text=element_text(size=14)) +
  scale_color_viridis_d(name = "Experience \nlevel", end = 0.7, option = "inferno") + 
  scale_fill_viridis_d(name = "Experience \nlevel", end =0.7, option = 'inferno') +
  theme(legend.position  = c(-1.0, -1.0), legend.direction = "horizontal", 
        legend.background = element_rect(fill="transparent", colour = NA),
        legend.title = element_text(size = 0), legend.text = element_text(size = 0),
        panel.spacing.x = unit(1, "lines"), 
        plot.margin = margin(c(0,0, 0, 0), unit = "cm"))
plot(Figure4a) #To see the figure in R

# Set date
DATE <- "4-28-2023"

tiff(filename=paste("Fig4a_Motor-Routines_", DATE, ".tif", sep=""), units="mm", width=150, height=200, res=1200)

plot(Figure4a) #To export it

setwd(PLOTDIR); 

dev.off()


#_______________________________________________________________________________________________________________________________#


# Question
#### Is the amount of pollen collected into the corbiculae per visit affected by plant species and treatment? ####

# Run the model
# Nested random effects; Bee within Colony
Pollen.Collected <- glmmTMB(log10(CorbPolperVis) ~ Treatment * Species + (1|ColonyID/BeeID), data = dataset1)

# Examine the model
summary(Pollen.Collected)

# Check Type II sums of squares, for Wald Tests
Anova(Pollen.Collected)

# Response: log10(CorbPolperVis)
#                        Chisq Df Pr(>Chisq)    
# Treatment          13.2649  1  0.0002704 ***
# Species           375.6822  3  < 2.2e-16 ***
# Treatment:Species   0.3865  3  0.9430084

emmeans(Pollen.Collected, list(pairwise ~ Treatment * Species), adjust = "tukey")

# Sjplot effect sizes, data, and predicted values
plot_model(Pollen.Collected)
plot_model(Pollen.Collected, type="pred", terms=c("Treatment"), show.data = T) #
plot_model(Pollen.Collected, type="pred", terms=c("Species", "Treatment"), show.data = T) #
plot_model(Pollen.Collected, type="pred", terms=c("TimeTotal", "Treatment"), show.data = T) #

# Model assumptions via DHARMa
simulationOutput1 <- simulateResiduals(fittedModel = Pollen.Collected, n = 10000)
plot(simulationOutput1) #Plot of QQ plot looks good; all residuals good but there is an outlier as of 7/16/2022 - so one of the new EXAF bees is responsible
testUniformity(simulationOutput = simulationOutput1) #Uniformity good 
testDispersion(simulationOutput1) #Dispersion good
testZeroInflation(simulationOutput1) #Zero inflation good


# Figure with paired data

# Rename the figure legends
dataset1$Treatment = plyr::mapvalues(dataset1$Treatment, from = c("DayOne", "DayTwo"), to = c("Naïve", "Experienced"))
dataset1$Species = plyr::mapvalues(dataset1$Species, from = c("EXAF", "MIGU", "PHCA", "SOEL"), 
                                 to = c("E. affine", "E. guttata", "P. campanularia", "S. elaeagnifolium"))

# Reorder the experience levels (treatment variables)
dataset1$Treatment <- factor(dataset1$Treatment, levels = c("Naïve","Experienced"))

# Plot figure
Figure5a <- ggplot(dataset1, aes(x = Treatment, y = log10(CorbPolperVis))) + 
  geom_boxplot(aes(fill = Treatment), alpha = .2) +
  geom_line(aes(group = BeeID)) + 
  geom_point(size = 1) + 
  facet_wrap(~ Species) +
  ylab("Log (pollen collected into pollen baskets per visit)") + xlab("Bee experience level") + theme_classic() + 
  theme(text=element_text(size=14)) +
  scale_color_viridis_d(name = "Experience \nlevel", end = 0.7, option = "inferno") + 
  scale_fill_viridis_d(name = "Experience \nlevel", end =0.7, option = 'inferno') +
  theme(legend.position  = c(-1.0, -1.0), legend.direction = "horizontal", 
        legend.background = element_rect(fill="transparent", colour = NA),
        legend.title = element_text(size = 0), legend.text = element_text(size = 0),
        panel.spacing.x = unit(1, "lines"), 
        plot.margin = margin(c(0,0, 0, 0), unit = "cm"))
plot(Figure5a) #To see the figure in R

# Set date
DATE <- "4-28-2023"

tiff(filename=paste("Fig5a_Corb-Pollen_", DATE, ".tif", sep=""), units="mm", width=120, height=200, res=1200)

plot(Figure5a) #To export it

setwd(PLOTDIR); 

dev.off()


#_______________________________________________________________________________________________________________________________#


# Question
#### Is the amount of pollen collected into the corbiculae per visit affected by plant species and treatment? ####

# Run the model
# Nested random effects; Bee within Colony
Pollen.Collected.Rate <- glmmTMB(log10(CorbPolperVisPerSec) ~ Treatment * Species + (1|ColonyID/BeeID), data = dataset1)

# Examine the model
summary(Pollen.Collected.Rate)

# Check Type II sums of squares, for Wald Tests
Anova(Pollen.Collected.Rate)

# Response: log10(CorbPolperVisPerSec)
#                     Chisq Df Pr(>Chisq)    
# Treatment           9.8808  1    0.00167 ** 
# Species           181.0640  3    < 2e-16 ***
# Treatment:Species   0.9233  3    0.81981

emmeans(Pollen.Collected.Rate, list(pairwise ~ Treatment * Species), adjust = "tukey")

# Sjplot effect sizes, data, and predicted values
plot_model(Pollen.Collected.Rate)
plot_model(Pollen.Collected.Rate, type="pred", terms=c("Treatment"), show.data = T) #
plot_model(Pollen.Collected.Rate, type="pred", terms=c("Species", "Treatment"), show.data = T) #
plot_model(Pollen.Collected, type="pred", terms=c("TimeTotal", "Treatment"), show.data = T) #

# Model assumptions via DHARMa
simulationOutput1 <- simulateResiduals(fittedModel = Pollen.Collected.Rate, n = 10000)
plot(simulationOutput1) #Plot of QQ plot looks good; all residuals good but there is an outlier as of 7/16/2022 - so one of the new EXAF bees is responsible
testUniformity(simulationOutput = simulationOutput1) #Uniformity good 
testDispersion(simulationOutput1) #Dispersion good
testZeroInflation(simulationOutput1) #Zero inflation good


# Figure with paired data

# Rename the figure legends
dataset1$Treatment = plyr::mapvalues(dataset1$Treatment, from = c("DayOne", "DayTwo"), to = c("Naïve", "Experienced"))
dataset1$Species = plyr::mapvalues(dataset1$Species, from = c("EXAF", "MIGU", "PHCA", "SOEL"), 
                                   to = c("E. affine", "E. guttata", "P. campanularia", "S. elaeagnifolium"))

# Reorder the experience levels (treatment variables)
dataset1$Treatment <- factor(dataset1$Treatment, levels = c("Naïve","Experienced"))

# Plot figure
FigureS1a <- ggplot(dataset1, aes(x = Treatment, y = log10(CorbPolperVisPerSec))) + 
  geom_boxplot(aes(fill = Treatment), alpha = .2) +
  geom_line(aes(group = BeeID)) + 
  geom_point(size = 1) + 
  facet_wrap(~ Species) +
  ylab("Log (pollen collection rate)") + xlab("Bee experience level") + theme_classic() + 
  theme(text=element_text(size=14)) +
  scale_color_viridis_d(name = "Experience \nlevel", end = 0.7, option = "inferno") + 
  scale_fill_viridis_d(name = "Experience \nlevel", end =0.7, option = 'inferno') +
  theme(legend.position  = c(-1.0, -1.0), legend.direction = "horizontal", 
        legend.background = element_rect(fill="transparent", colour = NA),
        legend.title = element_text(size = 0), legend.text = element_text(size = 0),
        panel.spacing.x = unit(1, "lines"), 
        plot.margin = margin(c(0,0, 0, 0), unit = "cm"))
plot(FigureS1a) #To see the figure in R

# Set date
DATE <- "6-7-2023"

tiff(filename=paste("FigS1a_Corb-Pollen-Rate_", DATE, ".tif", sep=""), units="mm", width=120, height=200, res=1200)

plot(FigureS1a) #To export it

setwd(PLOTDIR); 

dev.off()


#_______________________________________________________________________________________________________________________________#


# Question
#### Is the amount of pollen received by the stigma affected by plant species and treatment? ####


# Run the model
Pollen.Receipt <- glmmTMB(log10(StigPolperVis) ~ Treatment * Species + (1|ColonyID/BeeID), data = dataset1)

# Examine the model
summary(Pollen.Receipt)

# Check Type II sums of squares, for Wald Tests
Anova(Pollen.Receipt)

# Post-hoc test
emmeans(Pollen.Receipt, list(pairwise ~ Treatment * Species), adjust = "tukey")

# Response: log10(StigPolperVis)
#                   Chisq Df Pr(>Chisq)    
# Treatment           0.7569  1     0.3843    
# Species           128.6436  3     <2e-16 ***
# Treatment:Species   6.0705  3     0.1082


# Sjplot effect sizes, data, and predicted values
plot_model(Pollen.Receipt)
plot_model(Pollen.Receipt, type="pred", terms=c("Treatment"), show.data = T) #
plot_model(Pollen.Receipt, type="pred", terms=c("Species", "Treatment"), show.data = T) #
plot_model(Pollen.Receipt, type="pred", terms=c("TimeTotal", "Treatment"), show.data = T) #


# Model assumptions via DHARMa
simulationOutput1 <- simulateResiduals(fittedModel = Pollen.Receipt, n = 10000)
plot(simulationOutput1) #Plot of QQ plot looks good
testUniformity(simulationOutput = simulationOutput1) #Uniformity good 
testDispersion(simulationOutput1) #Dispersion good
testZeroInflation(simulationOutput1) #Zero inflation good


# Figure with paired data

# Rename the figure legends
dataset1$Treatment = plyr::mapvalues(dataset1$Treatment, from = c("DayOne", "DayTwo"), to = c("Naïve", "Experienced"))
dataset1$Species = plyr::mapvalues(dataset1$Species, from = c("EXAF", "MIGU", "PHCA", "SOEL"), 
                                   to = c("E. affine", "E. guttata", "P. campanularia", "S. elaeagnifolium"))

# Reorder the experience levels (treatment variables)
dataset1$Treatment <- factor(dataset1$Treatment, levels = c("Naïve","Experienced"))

# Plot figure
Figure6a <- ggplot(dataset1, aes(x = Treatment, y = log10(StigPolperVisPerSec))) + 
  geom_boxplot(aes(fill = Treatment), alpha = .2) +
  geom_line(aes(group = BeeID)) + 
  geom_point(size = 1) + 
  facet_wrap(~ Species) +
  ylab("Log (pollen deposited on the flower stigma per visit)") + xlab("Bee experience level") + theme_classic() + 
  theme(text=element_text(size=14)) +
  scale_color_viridis_d(name = "Experience \nlevel", end = 0.7, option = "inferno") + 
  scale_fill_viridis_d(name = "Experience \nlevel", end =0.7, option = 'inferno') +
  theme(legend.position  = c(-1.0, -1.0), legend.direction = "horizontal", 
        legend.background = element_rect(fill="transparent", colour = NA),
        legend.title = element_text(size = 0), legend.text = element_text(size = 0),
        panel.spacing.x = unit(1, "lines"), 
        plot.margin = margin(c(0,0, 0, 0), unit = "cm"))
plot(Figure6a) #To see the figure in R

# Set date
DATE <- "4-28-2023"

tiff(filename=paste("Fig6a_Pollen-Deposition_", DATE, ".tif", sep=""), units="mm", width=120, height=200, res=1200)

plot(Figure6a) #To export it

setwd(PLOTDIR); 

dev.off()


#_______________________________________________________________________________________________________________________________#


# Question
#### Is the amount of pollen received by the stigma affected by plant species and treatment? ####


# Run the model
Pollen.Receipt <- glmmTMB(log10(StigPolperVisPerSec) ~ Treatment * Species + (1|ColonyID/BeeID), data = dataset1)

# Examine the model
summary(Pollen.Receipt)

# Check Type II sums of squares, for Wald Tests
Anova(Pollen.Receipt)

# Post-hoc test
emmeans(Pollen.Receipt, list(pairwise ~ Treatment * Species), adjust = "tukey")

# Response: log10(StigPolperVisPerSec)
#                   Chisq Df Pr(>Chisq)    
# Treatment          0.5248  1     0.4688    
# Species           66.7475  3  2.121e-14 ***
# Treatment:Species  4.5423  3     0.2085


# Sjplot effect sizes, data, and predicted values
plot_model(Pollen.Receipt)
plot_model(Pollen.Receipt, type="pred", terms=c("Treatment"), show.data = T) #
plot_model(Pollen.Receipt, type="pred", terms=c("Species", "Treatment"), show.data = T) #
plot_model(Pollen.Receipt, type="pred", terms=c("TimeTotal", "Treatment"), show.data = T) #


# Model assumptions via DHARMa
simulationOutput1 <- simulateResiduals(fittedModel = Pollen.Receipt, n = 10000)
plot(simulationOutput1) #Plot of QQ plot looks good
testUniformity(simulationOutput = simulationOutput1) #Uniformity good 
testDispersion(simulationOutput1) #Dispersion good
testZeroInflation(simulationOutput1) #Zero inflation good


# Figure with paired data

# Rename the figure legends
dataset1$Treatment = plyr::mapvalues(dataset1$Treatment, from = c("DayOne", "DayTwo"), to = c("Naïve", "Experienced"))
dataset1$Species = plyr::mapvalues(dataset1$Species, from = c("EXAF", "MIGU", "PHCA", "SOEL"), 
                                   to = c("E. affine", "E. guttata", "P. campanularia", "S. elaeagnifolium"))

# Reorder the experience levels (treatment variables)
dataset1$Treatment <- factor(dataset1$Treatment, levels = c("Naïve","Experienced"))

# Plot figure
FigureS1b <- ggplot(dataset1, aes(x = Treatment, y = log10(StigPolperVisPerSec))) + 
  geom_boxplot(aes(fill = Treatment), alpha = .2) +
  geom_line(aes(group = BeeID)) + 
  geom_point(size = 1) + 
  facet_wrap(~ Species) +
  ylab("Log (pollen deposition rate)") + xlab("Bee experience level") + theme_classic() + 
  theme(text=element_text(size=14)) +
  scale_color_viridis_d(name = "Experience \nlevel", end = 0.7, option = "inferno") + 
  scale_fill_viridis_d(name = "Experience \nlevel", end =0.7, option = 'inferno') +
  theme(legend.position  = c(-1.0, -1.0), legend.direction = "horizontal", 
        legend.background = element_rect(fill="transparent", colour = NA),
        legend.title = element_text(size = 0), legend.text = element_text(size = 0),
        panel.spacing.x = unit(1, "lines"), 
        plot.margin = margin(c(0,0, 0, 0), unit = "cm"))
plot(FigureS1b) #To see the figure in R

# Set date
DATE <- "6-7-2023"

tiff(filename=paste("FigS1b_Pollen-Deposition-Rate_", DATE, ".tif", sep=""), units="mm", width=120, height=200, res=1200)

plot(FigureS1b) #To export it

setwd(PLOTDIR); 

dev.off()


#_______________________________________________________________________________________________________________________________#


# QUESTION 3
#### Does learning to successfully collect pollen differ among plant species?  ####

# Run the model
# Nested random effects; NumVisits within Bee
# QQ plot is not normal when ColonyID included
Handling.Effectiveness <- glmmTMB(log10(VisitsToCriterion) ~ Species * Treatment + (1 | ColonyID / BeeID), data = dataset1)

# Examine the model
summary(Handling.Effectiveness)

# Check Type II sums of squares, for Wald Tests
Anova(Handling.Effectiveness)

# Response: VisitsToCriterion
#                     Chisq Df Pr(>Chisq)    
# Species           58.8887  3  1.015e-12 ***
# Treatment          3.8818  1    0.04881 *  
# Species:Treatment 25.8747  3  1.013e-05 ***

# Post-hoc test
emmeans(Handling.Effectiveness, list(pairwise ~ Species * Treatment), adjust = "tukey")

# $`pairwise differences of Species`
# 1           estimate    SE   df t.ratio p.value
#  EXAF - MIGU  -11.019 1.65 138  -6.682  <.0001
# EXAF - PHCA    2.651 1.57 138   1.685  0.3353
# EXAF - SOEL    0.848 1.54 138   0.551  0.9462
# MIGU - PHCA   13.670 1.71 138   8.015  <.0001
# MIGU - SOEL   11.867 1.67 138   7.087  <.0001
# PHCA - SOEL   -1.803 1.60 138  -1.126  0.6740

# Model assumptions via DHARMa
simulationOutput1 <- simulateResiduals(fittedModel = Handling.Effectiveness, n = 1000)
plot(simulationOutput1) # QQ plot is good
testUniformity(simulationOutput = simulationOutput1) # Uniformity good
testDispersion(simulationOutput1) # Dispersion good
testZeroInflation(simulationOutput1) #Zero inflation good

# Model assumptions via sjPlot
# Normality of residuals and outliers
plot_model(Handling.Effectiveness, type = "diag") #


#_______________________________________________________________________________________________________________________________#


# Figure with paired data

# Rename the figure legends
dataset1$Treatment = plyr::mapvalues(dataset1$Treatment, from = c("DayOne", "DayTwo"), to = c("Naïve", "Experienced"))
dataset1$Species = plyr::mapvalues(dataset1$Species, from = c("EXAF", "MIGU", "PHCA", "SOEL"), 
                                   to = c("E. affine", "E. guttata", "P. campanularia", "S. elaeagnifolium"))

# Reorder the experience levels (treatment variables)
dataset1$Treatment <- factor(dataset1$Treatment, levels = c("Naïve","Experienced"))

# Plot figure
Figure6a <- ggplot(dataset1, aes(x = Treatment, y = log10(VisitsToCriterion))) + 
  geom_boxplot(aes(fill = Treatment), alpha = .2) +
  geom_line(aes(group = BeeID)) + 
  geom_point(size = 1) + 
  facet_wrap(~ Species) +
  ylab("Log (number of visits until learning criterion met)") + xlab("Bee experience level") + theme_classic() + 
  theme(text=element_text(size=14)) +
  scale_color_viridis_d(name = "Experience \nlevel", end = 0.7, option = "inferno") + 
  scale_fill_viridis_d(name = "Experience \nlevel", end =0.7, option = 'inferno') +
  theme(legend.position  = c(-1.0, -1.0), legend.direction = "horizontal", 
        legend.background = element_rect(fill="transparent", colour = NA),
        legend.title = element_text(size = 0), legend.text = element_text(size = 0),
        panel.spacing.x = unit(1, "lines"), 
        plot.margin = margin(c(0,0, 0, 0), unit = "cm"))
plot(Figure6a) #To see the figure in R

# Set date
DATE <- "5-29-2023"

tiff(filename=paste("Fig6a_Handling-Effectiveness_", DATE, ".tif", sep=""), units="mm", width=120, height=200, res=1200)

plot(Figure6a) #To export it

setwd(PLOTDIR); 

dev.off()


#_______________________________________________________________________________________________________________________________#


# QUESTION 4
#### Does experience affect bees making errors on handling flowers just on their very first visit? ####
# Number of naive, experienced bees that failed to collect from anthers on first visit

# 2-sample test for equality of proportions with continuity correction
# EXAF bees
prop.test(x = c(14, 17), n = c(21, 21), alternative = "two.sided", correct = TRUE)

# Results
# X-squared = 0.49267, df = 1, p-value = 0.4827


# 2-sample test for equality of proportions with continuity correction
# MIGU bees
prop.test(x = c(9, 5), n = c(20, 20), alternative = "two.sided", correct = TRUE)

# Results
# X-squared = 0.98901, df = 1, p-value = 0.32


# 2-sample test for equality of proportions with continuity correction
# PHCA bees
prop.test(x = c(13, 17), n = c(17, 19), alternative = "two.sided", correct = TRUE)

# Results
# X-squared = 0.35666, df = 1, p-value = 0.5504


# 2-sample test for equality of proportions with continuity correction
# SOEL bees
prop.test(x = c(20, 19), n = c(20, 20), alternative = "two.sided", correct = TRUE)

# Results
# X-squared = 7.8886e-31, df = 1, p-value = 1


# 2-sample test for equality of proportions with continuity correction
# Naive bees, in order: EXAF, MIGU, PHCA, SOEL
prop.test(x = c(14, 9, 13, 20), n = c(21, 20, 17, 20), alternative = "two.sided", correct = TRUE)

# Results
# X-squared = 15.404, df = 3, p-value = 0.001502


#_______________________________________________________________________________________________________________________________#

#### Handling time traces ####

#### Summarizing data function ####
# Gives count, mean, standard deviation, standard error of the mean, and confidence interval (default 95%).
#   data: a data frame.
#   measurevar: the name of a column that contains the variable to be summariezed
#   groupvars: a vector containing names of columns that contain grouping variables
#   na.rm: a boolean that indicates whether to ignore NA's
#   conf.interval: the percent range of the confidence interval (default is 95%)

summarySE <- function(data=NULL, measurevar, groupvars=NULL, na.rm=FALSE,
                      conf.interval=.95, .drop=TRUE) {
  library(plyr)
  
  # New version of length which can handle NA's: if na.rm==T, don't count them
  length2 <- function (x, na.rm=FALSE) {
    if (na.rm) sum(!is.na(x))
    else       length(x)
  }
  
  # This does the summary. For each group's data frame, return a vector with
  # N, mean, and sd
  datac <- ddply(data, groupvars, .drop=.drop,
                 .fun = function(xx, col) {
                   c(N    = length2(xx[[col]], na.rm=na.rm),
                     mean = mean   (xx[[col]], na.rm=na.rm),
                     sd   = sd     (xx[[col]], na.rm=na.rm)
                   )
                 },
                 measurevar
  )
  
  # Rename the "mean" column    
  datac <- rename(datac, c("mean" = measurevar))
  
  datac$se <- datac$sd / sqrt(datac$N)  # Calculate standard error of the mean
  
  # Confidence interval multiplier for standard error
  # Calculate t-statistic for confidence interval: 
  # e.g., if conf.interval is .95, use .975 (above/below), and use df=N-1
  ciMult <- qt(conf.interval/2 + .5, datac$N-1)
  datac$ci <- datac$se * ciMult
  
  return(datac)
}


# Find the summary of the data
dataset2 <- summarySE(dataset2, measurevar="HandlingTime", groupvars=c("Treatment", "VisitNo"), na.rm=TRUE)

dataset3 <- summarySE(dataset3, measurevar="HandlingTime", groupvars=c("Treatment", "VisitNo"), na.rm=TRUE)

dataset4 <- summarySE(dataset4, measurevar="HandlingTime", groupvars=c("Treatment", "VisitNo"), na.rm=TRUE)

dataset5 <- summarySE(dataset5, measurevar="HandlingTime", groupvars=c("Treatment", "VisitNo"), na.rm=TRUE)

# The errorbars overlapped, so use position_dodge to move them horizontally
pd <- position_dodge(0.1) # move them .05 to the left and right


#### Figure for PHCA handling time trace ####

# Rename the figure legends
dataset2$title <- "P. campanularia"
dataset2$Treatment = plyr::mapvalues(dataset2$Treatment, from = c("DayOne", "DayTwo"), to = c("Naïve", "Experienced"))

# Reorder the experience levels (treatment variables)
dataset2$Treatment <- factor(dataset2$Treatment, levels = c("Naïve","Experienced"))

# Black error bars - notice the mapping of 'group=Treatment' -- without it, the error bars won't be dodged!
Figure1a <- ggplot(dataset2, aes(x=VisitNo, y=HandlingTime, colour=Treatment, group=Treatment)) + 
  geom_errorbar(aes(ymin=HandlingTime-se, ymax=HandlingTime+se), colour="black", width=.1, position=pd) +
  geom_line(position=pd) +
  geom_point(position=pd, size=2, shape=21, fill="white") + # 21 is filled circle
  xlab("Visit number") + ylab("Flower handling time (s) ± SE") + theme_classic() + 
  theme(text=element_text(size=14)) +
  scale_color_viridis_d(name = "Experience \nlevel", end = 0.7, option = "inferno") + 
  scale_fill_viridis_d(name = "Experience \nlevel", end =0.7, option = 'inferno') +
  guides(colour=guide_legend(nrow=2, byrow=TRUE)) +  theme_classic() + 
  theme(legend.position  =c(0.5, 0.5), legend.direction = "horizontal", 
        legend.title = element_text(size = 12), legend.text = element_text(size = 12)) + 
  facet_grid(. ~ title) + theme(strip.text.x = element_text(size = 12)) +
  xlim(0,40) + ylim(0,105)
plot(Figure1a) #To see the figure in R


# Set date
DATE <- "5-24-2023"

tiff(filename=paste("Fig1a_Handling-Time-Trace-PHCA_", DATE, ".tif", sep=""), units="mm", width=120, height=150, res=1200)

plot(Figure1a) #To export it

setwd(PLOTDIR); 

dev.off()


#### Figure for EXAF handling time trace ####

# Rename the figure legends
dataset3$title <- "E. affine"
dataset3$Treatment = plyr::mapvalues(dataset3$Treatment, from = c("DayOne", "DayTwo"), to = c("Naïve", "Experienced"))

# Reorder the experience levels (treatment variables)
dataset3$Treatment <- factor(dataset3$Treatment, levels = c("Naïve","Experienced"))

# Black error bars - notice the mapping of 'group=Treatment' -- without it, the error bars won't be dodged!
Figure1b <- ggplot(dataset3, aes(x=VisitNo, y=HandlingTime, colour=Treatment, group=Treatment)) + 
  geom_errorbar(aes(ymin=HandlingTime-se, ymax=HandlingTime+se), colour="black", width=.1, position=pd) +
  geom_line(position=pd) +
  geom_point(position=pd, size=2, shape=21, fill="white") + # 21 is filled circle
  xlab("Visit number") + ylab("Flower handling time (s) ± SE") + theme_classic() + 
  theme(text=element_text(size=14)) +
  scale_color_viridis_d(name = "Experience \nlevel", end = 0.7, option = "inferno") + 
  scale_fill_viridis_d(name = "Experience \nlevel", end =0.7, option = 'inferno') +
  guides(colour=guide_legend(nrow=2, byrow=TRUE)) +  theme_classic() + 
  theme(legend.position  =c(0.5, 0.5), legend.direction = "horizontal", 
        legend.title = element_text(size = 12), legend.text = element_text(size = 12)) + 
  facet_grid(. ~ title) + theme(strip.text.x = element_text(size = 12)) +
  xlim(0,40) + ylim(0,105)
plot(Figure1b) #To see the figure in R

# Set date
DATE <- "5-24-2023"

tiff(filename=paste("Fig1b_Handling-Time-Trace-EXAF_", DATE, ".tif", sep=""), units="mm", width=120, height=150, res=1200)

plot(Figure1b) #To export it

setwd(PLOTDIR); 

dev.off()


#### Figure for SOEL handling time trace ####

# Rename the figure legends
dataset4$title <- "S. elaeagnifolium"
dataset4$Treatment = plyr::mapvalues(dataset4$Treatment, from = c("DayOne", "DayTwo"), to = c("Naïve", "Experienced"))

# Reorder the experience levels (treatment variables)
dataset4$Treatment <- factor(dataset4$Treatment, levels = c("Naïve","Experienced"))

# Black error bars - notice the mapping of 'group=Treatment' -- without it, the error bars won't be dodged!
Figure1c <- ggplot(dataset4, aes(x=VisitNo, y=HandlingTime, colour=Treatment, group=Treatment)) + 
  geom_errorbar(aes(ymin=HandlingTime-se, ymax=HandlingTime+se), colour="black", width=.1, position=pd) +
  geom_line(position=pd) +
  geom_point(position=pd, size=2, shape=21, fill="white") + # 21 is filled circle
  xlab("Visit number") + ylab("Flower handling time (s) ± SE") + theme_classic() + 
  theme(text=element_text(size=14)) +
  scale_color_viridis_d(name = "Experience \nlevel", end = 0.7, option = "inferno") + 
  scale_fill_viridis_d(name = "Experience \nlevel", end =0.7, option = 'inferno') +
  guides(colour=guide_legend(nrow=2, byrow=TRUE)) +  theme_classic() + 
  theme(legend.position  =c(0.5, 0.5), legend.direction = "horizontal", 
        legend.title = element_text(size = 12), legend.text = element_text(size = 12)) + 
  facet_grid(. ~ title) + theme(strip.text.x = element_text(size = 12)) +
  xlim(0,40) + ylim(0,105)
plot(Figure1c) #To see the figure in R

# Set date
DATE <- "5-24-2023"

tiff(filename=paste("Fig1c_Handling-Time-Trace-SOEL_", DATE, ".tif", sep=""), units="mm", width=120, height=150, res=1200)

plot(Figure1c) #To export it

setwd(PLOTDIR); 

dev.off()


#### Figure for MIGU handling time trace ####

# Rename the figure legends
dataset5$title <- "E. guttata"
dataset5$Treatment = plyr::mapvalues(dataset5$Treatment, from = c("DayOne", "DayTwo"), to = c("Naïve", "Experienced"))

# Reorder the experience levels (treatment variables)
dataset5$Treatment <- factor(dataset5$Treatment, levels = c("Naïve","Experienced"))

# Black error bars - notice the mapping of 'group=Treatment' -- without it, the error bars won't be dodged!
Figure1d <- ggplot(dataset5, aes(x=VisitNo, y=HandlingTime, colour=Treatment, group=Treatment, cex.main=3.5)) + 
  geom_errorbar(aes(ymin=HandlingTime-se, ymax=HandlingTime+se), colour="black", width=.1, position=pd) +
  geom_line(position=pd) +
  geom_point(position=pd, size=2, shape=21, fill="white") + # 21 is filled circle
  xlab("Visit number") + ylab("Flower handling time (s) ± SE") + theme_classic() + 
  theme(text=element_text(size=14)) +
  scale_color_viridis_d(name = "Experience \nlevel", end = 0.7, option = "inferno") + 
  scale_fill_viridis_d(name = "Experience \nlevel", end =0.7, option = 'inferno') +
  guides(colour=guide_legend(nrow=2, byrow=TRUE)) +  theme_classic() + 
  theme(legend.position  =c(0.5, 0.5), legend.direction = "horizontal", 
        legend.title = element_text(size = 12), legend.text = element_text(size = 12)) + 
  facet_grid(. ~ title) + theme(strip.text.x = element_text(size = 12)) +
  xlim(0,40) + ylim(0,105)
plot(Figure1d) #To see the figure in R

# Set date
DATE <- "5-24-2023"

tiff(filename=paste("Fig1d_Handling-Time-Trace-MIGU_", DATE, ".tif", sep=""), units="mm", width=120, height=150, res=1200)

plot(Figure1d) #To export it

setwd(PLOTDIR); 

dev.off()
```
